# Altered striatal dopamine regulation in *ADGRL3* knockout mice

**DOI:** 10.1101/2025.07.31.667389

**Authors:** Nicole A. Perry-Hauser, Arturo Torres-Herraez, Siham Boumhaouad, Emily A. Makowicz, Daniel C. Lowes, Michelle Jin, Christine A. Denny, David Sulzer, Eugene V. Mosharov, Christoph Kellendonk, Jonathan A. Javitch

## Abstract

Dopaminergic signaling is essential for regulating movement, learning, and reward. Disruptions in this system are linked to neuropsychiatric disorders such as ADHD. ADGRL3, an adhesion G protein-coupled receptor highly expressed in the brain, is genetically associated with increased ADHD risk. *ADGRL3* knockout in animals alters expression of dopaminergic markers and induces dopamine-related behavioral changes. However, its precise role in modulating dopamine signaling remains unclear. We investigated how *ADGRL3* knockout affects striatal dopamine release in mice using *ex vivo* fast-scan cyclic voltammetry and *in vivo* fiber photometry with a dopamine sensor. *Ex vivo* measurements showed increased electrically-evoked dopamine release across the striatum. Conversely, *in vivo* recordings revealed reduced task-induced dopamine signals in the nucleus accumbens during an operant fixed interval task. This reduction was not due to impaired dopamine availability, as amphetamine-evoked release was unchanged. These findings suggest ADGRL3 modulates dopamine release in complex ways via different pre- and postsynaptic mechanisms.

## Introduction

*ADGRL3* (also known as *CL3*, *LEC3*, *CIRL3*, or *LPHN3*) encodes an adhesion G protein-coupled receptor (aGPCR) belonging to the latrophilin subfamily. It is predominantly expressed in the nervous system, where its levels peak shortly after birth and gradually decrease to adult levels during maturation (*1*). Single-nucleotide polymorphisms in *ADGRL3* that result in haploinsufficiency have been associated with an increased risk of attention-deficit/hyperactivity disorder (ADHD) and are predictive of susceptibility to substance use disorders (*1*, *2*). These associations suggest a crucial role for ADGRL3 in neurodevelopment. Consistent with this, ADGRL3 is thought to regulate hippocampal synapse formation, primarily through trans-synaptic complexes with endogenous ligands such as teneurins and fibronectin leucine-rich repeat transmembrane proteins (*3–9*).

Imbalances in the dopamine system are hypothesized to contribute to the pathophysiology of ADHD (*10*), and knockdown of *ADGRL3* in fish, as well as knockout in flies, mice, and rats, disrupts normal dopamine signaling and leads to abnormalities in dopamine-related behaviors, most notably increased locomotor activity (*11–17*). Initial characterization of an *ADGRL3* knockout (KO) mouse line revealed elevated whole brain mRNA expression of the dopamine transporter gene (*Dat1*), dopamine receptor D4 (*Drd4*), and tyrosine hydroxylase (*Th*) at postnatal day 0 (P0) (*15*). Neurochemical analysis further demonstrated significantly higher dopamine levels in the dorsal striatum of these mice, a brain region that receives dense dopaminergic innervation. In contrast, a constitutive *ADGRL3* KO rat model did not exhibit changes in striatal dopamine, norepinephrine, serotonin, or their metabolites at the biochemical level (*16*). However, several striatal dopamine-related proteins showed altered expression: tyrosine hydroxylase (TH), aromatic L-amino acid decarboxylase (AADC), and dopamine transporter (DAT) levels were increased, while dopamine receptor D1 and dopamine-regulated neuronal phosphoprotein (DARPP-32) were downregulated. Collectively, these findings suggest a role for ADGRL3 in regulating striatal dopamine neurotransmission, but the mechanisms underlying this regulation remain unclear.

Dopamine plays a critical role in various behaviors, including movement, learning, and motivation (*18*, *19*). Previous studies have assessed behavioral disturbances associated with dopamine dysfunction in *ADGRL3* KO mice (*13–15*). Motor control has been evaluated using the rotarod test, open field test, and gait analysis, revealing hyperactivity and unstable footing for *ADGRL3* KO animals. Learning and memory have been assessed through the delayed response task, novel object recognition task, and Barnes maze, where the *ADGRL3* KO mice exhibited significant deficits. Motivation and reward-seeking behaviors have been examined using a fixed ratio schedule of reinforcement, the forced swim test, and the continuous performance test. These results were more variable, suggesting that motivation depended on the specific task and reward context. Overall, *ADGRL3* KO mice exhibit disturbances in multiple dopamine-related behaviors, and the reported differences in dopamine signaling underscore the need for further investigation.

Here, we combined behavioral assays with dopamine measurements to investigate how loss of *ADGRL3* alters striatal function. We first validated locomotor phenotypes in *ADGRL3* KO mice using both open field and home-cage environments. To assess striatal dopamine signaling, we used two complementary approaches. First, we used *ex vivo* fast-scan cyclic voltammetry (FSCV) in acute striatal slices to measure electrically evoked dopamine release. Then, to examine dopamine dynamics in freely behaving animals, we used *in vivo* fiber photometry with the genetically encoded dopamine sensor dLight1.2 while mice performed an operant task.

Behavioral testing included a continuous reinforcement task followed by a fixed interval reinforcement schedule. Finally, we tested the capacity for dopamine release *in vivo* using an amphetamine challenge coupled to fiber photometry. Collectively, our findings suggest that ADGRL3 has a complex effect on striatal dopamine function, likely involving both pre- and postsynaptic mechanisms.

## Results

### Increased horizontal locomotion and decreased rearing behavior in *ADGRL3* KO mice across development

We assessed locomotor activity in WT and *ADGRL3* KO mice using a standard open field test. In this test, mice were placed in a plexiglass enclosure for 60 min undisturbed (**Fig. 1, Figs. S1** and **S2**). Prior to testing, mice were weighed, and a two-way mixed ANOVA was performed to analyze the effect of age and genotype on weight (**Fig. 1A, Fig. S1A, E**). *ADGRL3* KO mice had a statistically significant decrease in weight across development compared to WT controls.

**Figure 1.**
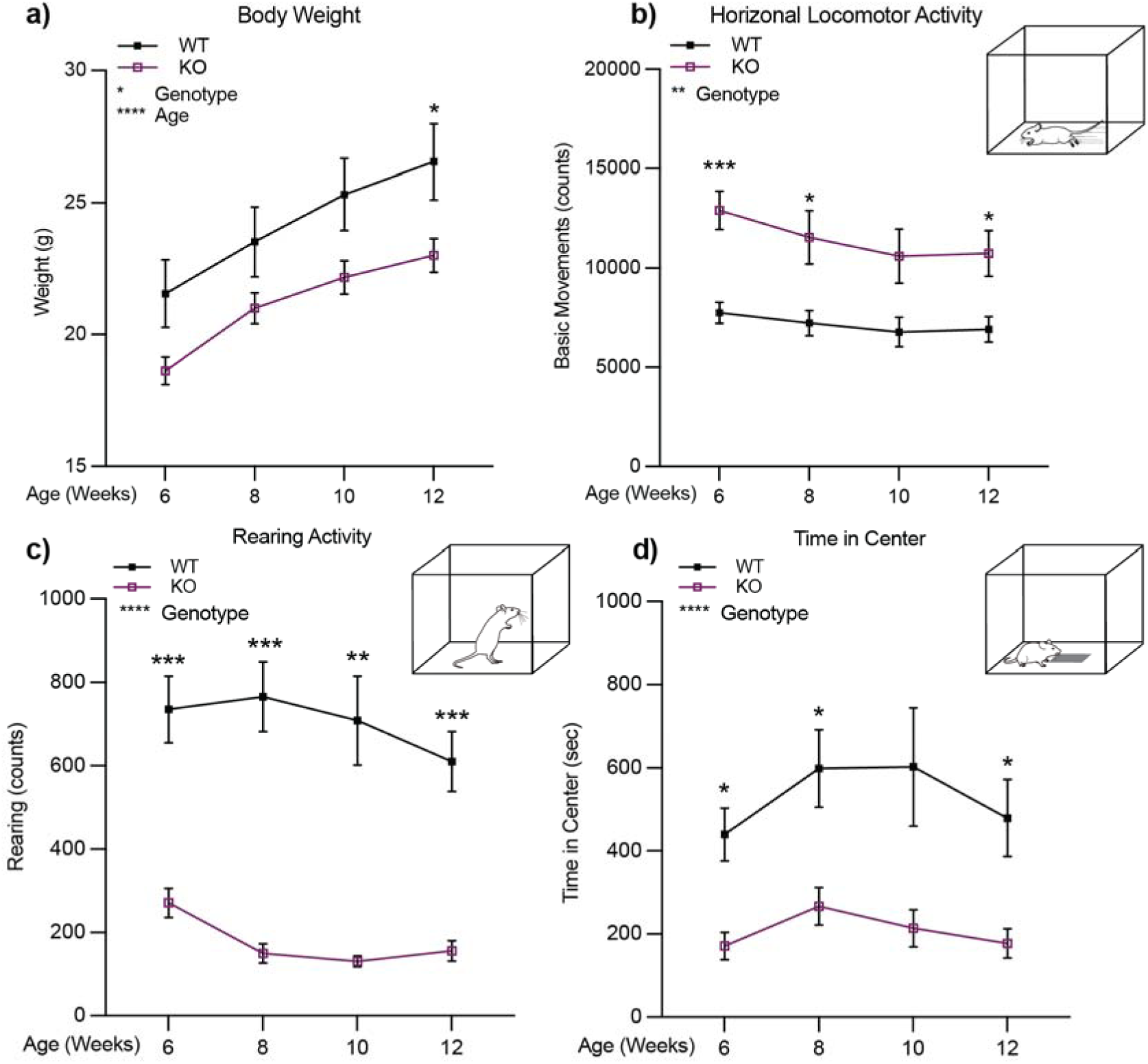
*ADGRL3* knockout mice show increased horizontal locomotion and decreased rearing activity across development. **(a)** Weight of wild-type (WT) and *ADGRL3* knockout (KO) mice. (**b)** Horizontal locomotor activity, **(c)** rearing activity, and **(d)** time in center for WT and *ADGRL3* KO mice across development (N=9 WT, 17 KO). A two-way repeated measures ANOVA was performed to analyze the effect of age and genotype on each dependent variable (weight, basic movements, rearing, or time in center). This was followed by ídák’s multiple comparisons test (*, p < 0.05; **, p<0.01; ***, p<0.001; ****, p < 0.0001).

*ADGRL3* KO mice displayed greater horizontal locomotor activity than WT mice (**Fig. 1B, Fig. S1B, F**). This increase in locomotor activity persisted throughout the 12-week testing period and into adulthood. This was further supported by independent tracking of X- and Y-axis ambulation (**Fig. S2A-B**), which showed that *ADGRL3* KO mice consistently traveled greater distances in both axes across all time points. These findings align with previous studies reporting hyperactivity in *ADGRL3* KO mice compared to heterozygous and WT littermates (*13*, *15*).

*ADGRL3* KO mice also showed a significant decrease in the time spent rearing (**Fig. 1C, Fig. S1C, G**), consistent with previous findings (*13*). However, unlike in previous reports, *ADGRL3* KO mice spent significantly less time in the center of the enclosure (**Fig. 1D, Fig. S1D, H**). These results were further corroborated by a separate analysis of spatial distribution in the open field, which showed that *ADGRL3* KO mice spent less time in the center and significantly more time in the periphery compared to WT controls (**Fig. S2C-D**). While both rearing and center time are frequently used as indicators of anxiety-like behavior (*20*), it is difficult to rule out the confounding influence of hyperactivity on these measures.

### *ADGRL3* knockout mice exhibit increased activity during the dark cycle compared to controls

To determine whether the increased locomotor activity observed for *ADGRL3* KO mice is time-of-day dependent and to assess potential effects on their sleep/wake cycles, we next monitored their behavior using the PiezoSleep System. WT and *ADGRL3* KO mice were individually housed for five consecutive days with *ad libitum* access to food and water under a 12-hour light/dark cycle. Floor sensors continuously tracked movement and sleep bouts. We plotted activity data across zeitgeber time (**Fig. 2A**) and found that *ADGRL3* KO mice had increased activity between light and dark cycles compared to WT controls (**Fig. 2B**). This heightened activity could be attributed to pronounced hyperactivity during the dark (active) cycle where *ADGRL3* KO mice spent significantly less time asleep with shorter sleep bouts (**Figs. 2C, D**). These findings align with previous reports in *ADGRL3* KO rats, where hyperactivity was more pronounced in the dark cycle (*16*).

**Figure 2.**
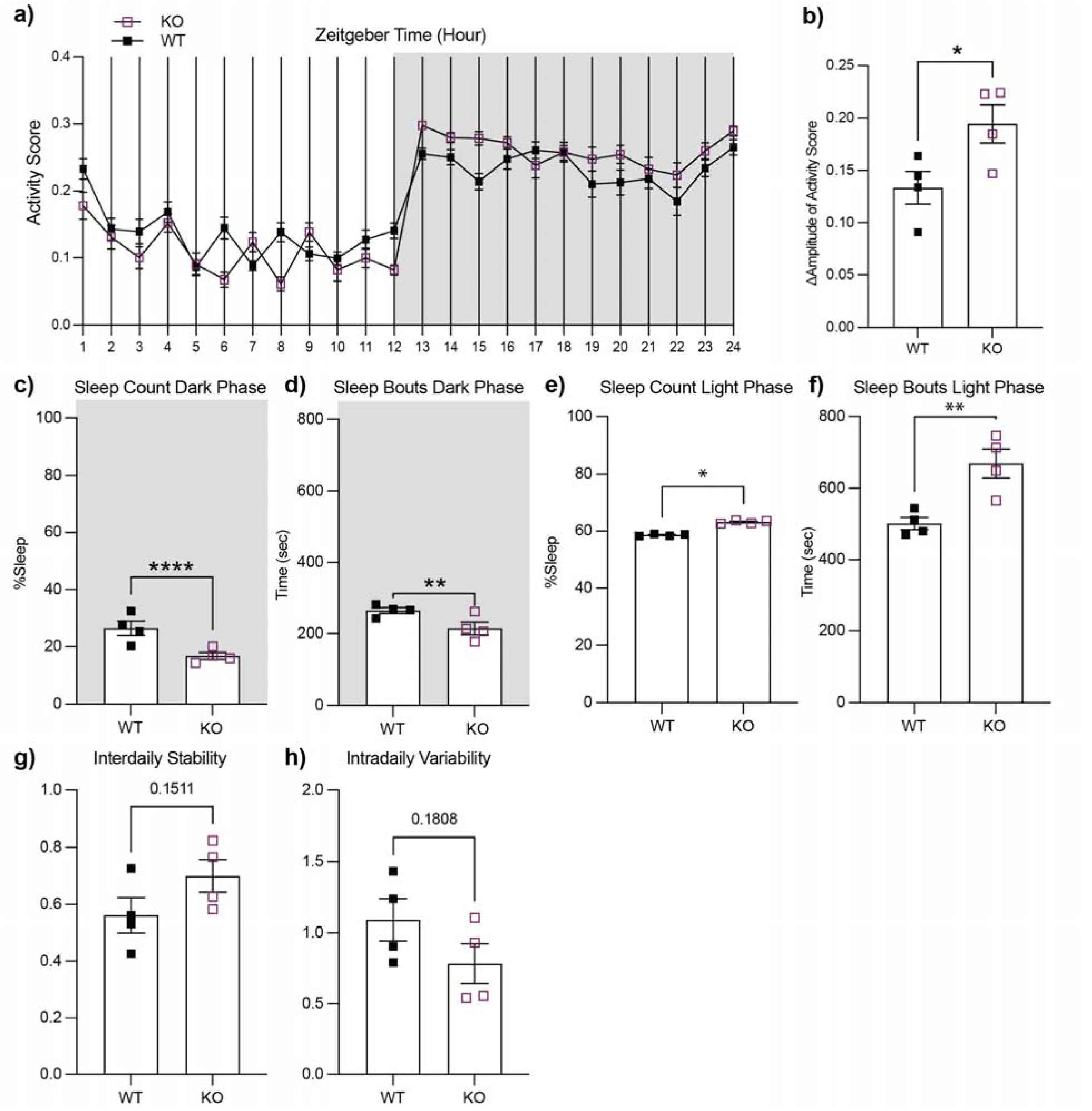
*ADGRL3* knockout mice exhibit increased activity in PiezoSleep boxes during the dark cycle compared to controls. Activity levels and sleep/wake cycles in wild-type (WT) and *ADGRL3* knockout (KO) mice were monitored for five consecutive days using the Piezo-Sleep mouse behavioral tracking system (Signal Solutions LLC). **(a)** Average activity scores for WT and *ADGRL3* KO mice presented in Zeitgeber time (N=4 WT, 4 KO). **(b)** Average change in the amplitude of activity for WT and *ADGRL3* KO mice. **(c)** Percentage of time spent asleep and **(d)** average duration of sleep bouts during the dark cycle. **(e)** Percentage of time spent asleep and **(f)** average duration of sleep bouts during the light cycle. **(g)** Interdaily stability was calculated to quantify the rest-activity rhythms for each mouse between different days. A value closer to 1 indicates stronger coupling to a zeitgeber with a period length of 24 hours (*50*). **(h)** Intradaily variability was calculated to quantify the fragmentation of the rest-activity pattern. This value converges to 0 for a perfect sine wave and approaches two for Gaussian noise (*50*). All statistical analyses were conducted using an Unpaired t-test (****, p < 0.0001; ***, p < 0.001; **, p < 0.01; and *, p < 0.05).

During the light cycle, *ADGRL3* KO mice spent significantly more time asleep with longer sleep bouts (**Fig. 2E, F).** Given their overall heightened activity compared to WT mice, this may indicate that their hyperactivity is particularly sensitive to light cues and/or circadian timing. To explore this further, we analyzed interdaily stability, a measure of how consistent daily activity patterns are, and intradaily variability, which reflects the fragmentation of activity within a 24-hour period (**Fig. 2G, H).** Although differences were not statistically significant, *ADGRL3* KO mice showed a trend toward higher interdaily stability and lower intradaily variability, suggesting a more robust and consolidated circadian rhythm regulated by external cues.

### *ADGRL3* KO mice show increased lick responses for an evaporated milk reward compared to WT mice

To assess hedonic drive in *ADGRL3* KO mice relative to WT controls, we used a “Davis Rig” gustometer, an apparatus designed for micro-analysis of ingestive behavior toward liquids in rodents (**Fig. 3**) (*21*). This system is commonly used prior to operant testing to evaluate unconditioned licking responses and detect baseline differences in consumption across experimental groups. While previous studies in *ADGRL3* KO mice have investigated motivational response to food rewards, the hedonic component of that motivation (i.e., the “liking” aspect) has not been directly addressed.

**Figure 3.**
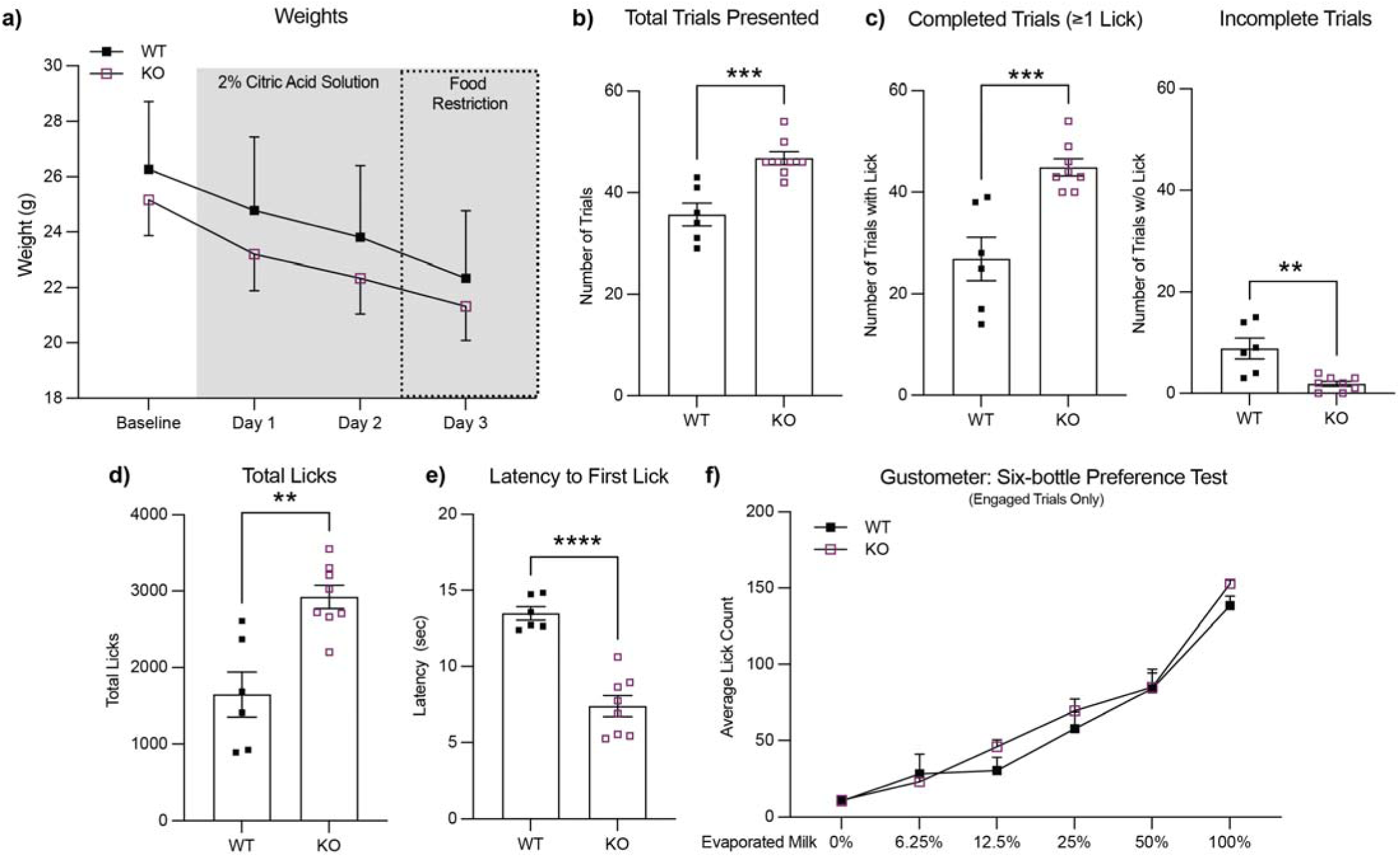
*ADGRL3* knockout mice show increased lick responses but similar taste preference for an evaporated milk reward compared to WT mice. Taste preference for and licking behavior toward an evaporated milk reward were measured for WT and *ADGRL3* knockout (KO) mice using a “Davis Rig” gustometer (N=6 WT, 8 KO). **(a)** Average weights for each genotype over the course of the study. **(b)** Number of trials presented, **(c)** number of completed and incomplete trials, **(d)** and total number of licks across the 30 min session. Statistical analysis consisted of an unpaired t test (***, p<0.001; **, p<0.01). **(e)** Average latency until first lick by genotype. Statistical analysis consisted of an unpaired t test (****, p<0.0001). **(f)** Average licks by each genotype for 6 different concentrations of evaporated milk (0%, 6.25%, 12.5%, 25%, 50%, and 100%). Statistical analysis consisted of a Two-way ANOVA followed by ídák’s multiple comparison test (****, p<0.0001 for % of evaporated milk). No statistical difference was detected for genotype or reward.

To this end, we conducted a six-bottle preference test, presenting water and five concentrations of evaporated milk (6.25%, 12.5%, 25%, 50%, and 100%) in pseudorandom order. The task lasted for 30 minutes, with a 20-second intertrial interval. Thus, mice that initiated licking more quickly and consistently could complete more trials within the fixed session duration. *ADGRL3* KO mice had a greater number of total trials and had more completed trials, defined as any trial in which at least one lick was made, compared to WT mice (**Fig. 3B, C**). They also licked more compared to WT controls and had a shorter latency to the first lick (**Fig. 3D, E**). Together, these findings suggest that *ADGRL3* KO mice have an increased ‘liking’ for the evaporated milk reward (*22*).

When analysis was restricted to completed trials—those in which mice initiated licking—the average lick count per solution did not differ between genotypes (**Fig. 3F Fig. S3**). This finding suggests that the increased task performance in *ADGRL3* KO mice may not be solely attributable to increased “liking” per se but rather reflect increased instrumental responding, a greater tendency to initiate and perform goal-directed actions to obtain a reward.

### *ADGRL3* KO mice have greater levels of evoked dopamine release in the striatum

Fast-scan cyclic voltammetry (FSCV) is a high-resolution electrochemical technique used to detect neurotransmitters with sub-second precision (*23*). This method relies on microelectrodes, with carbon-fiber microelectrodes being particularly well-suited for detecting cations like dopamine. We measured electrically evoked dopamine release *ex vivo* in acute brain slices containing the dorsal and ventral striatum from WT and *ADGRL3* KO mice aged 3-6 months (**Fig. 4**). Compared to WT controls, *ADGRL3* KO mice showed significantly higher levels of evoked dopamine release in both the dorsal (**Fig. 4B, D**) and ventral striatum (**Fig. 4F, H**). Despite the elevated response, there were no significant genotype differences in dopamine reuptake in either region, as indicated by the rate of dopamine signal decay (Tau1) following stimulation (**Fig. 4C, G**).

**Figure 4.**
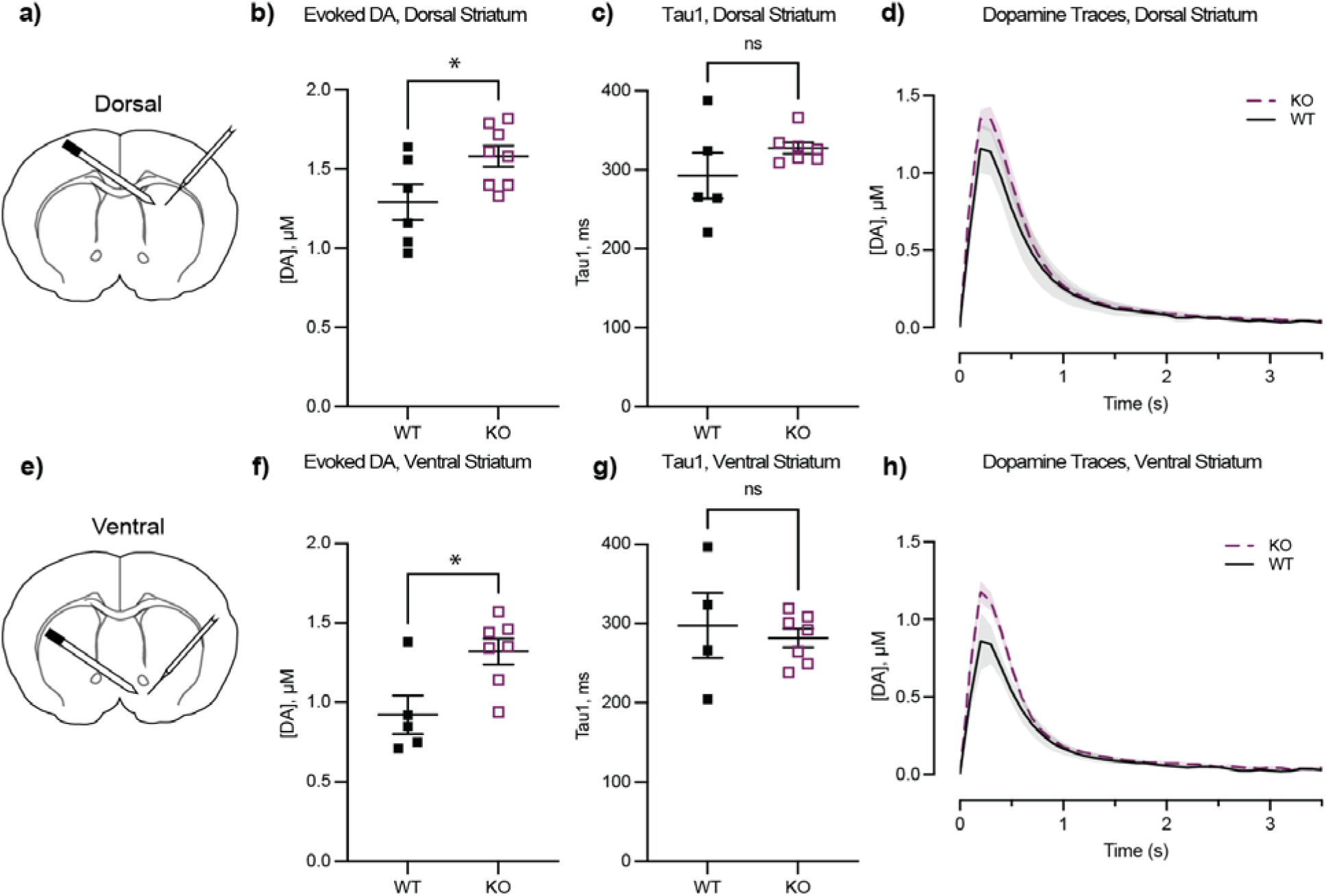
*ADGRL3* KO mice have greater levels of evoked dopamine (DA) and slower DA reuptake in the striatum. Coronal sections that included striatum were obtained from wild-type (WT) and *ADGRL3* knockout (KO) mice aged P102-P183. **(a)** Schematic of location for fast-scan cyclic voltammetry (FSCV) recordings from the dorsal striatum. Slices were stimulated with a bipolar concentric electrode and DA was detected using a carbon fiber electrode. **(b)** FSCV recordings of evoked DA release from the dorsal striatum (N=6 WT, N=8 KO; 31 slices WT, 44 slices KO). **(c)** DA reuptake from the dorsal striatum (N=5 WT, N=7 KO; 25 slices WT, 47 slices KO). **(d)** Averaged dopamine transients in response to a single electrical pulse in the dorsal striatum from all tested mice, illustrating signal kinetics. **(e)** Schematic of location for FSCV recordings from the ventral striatum. **(f)** FSCV recordings of evoked DA release from the ventral striatum (N=5 WT, N=7 KO; 24 slices WT, 36 slices KO). **(g)** DA reuptake from the ventral striatum (N=4 WT, N=7 KO; 19 slices WT, 42 slices KO). **(h)** Averaged dopamine transients in response to a single electrical pulse in the ventral striatum from all tested mice, illustrating signal kinetics. All quantal release events were analyzed in IgorPro using a publicly available GitHub-based analysis program (*53*) Peak parameters included amplitude (Imax, pA) and event duration (Tau1, ms). All statistical analyses were performed using an unpaired t-test (*, p<0.05).

### *ADGRL3* KO mice have similar levels of dopamine release in a continuous reinforcement task

In the nucleus accumbens, dopamine is released in response to unexpected rewards (*19*, *24*). However, with continued training, as rewards become predictable, dopamine release in response to rewards decreases and shifts to environmental cues that signal the rewards. Based on prior studies (*17*) and our *ex vivo* FSCV data showing differences in the releasable pool of dopamine, we hypothesized that *ADGRL3* KO mice would have greater cue-induced dopamine release in the striatum compared to WT controls. To test this hypothesis, we used *in vivo* fiber photometry with the genetically encoded fluorescent biosensor, dLight1.2 (*25*), to measure dopamine levels during a continuous reinforcement task and a fixed interval task.

Four weeks after viral injection and optic fiber implantation, mice began operant training. The initial phase, trough training, involved teaching the mice to access an evaporated milk reward delivered through a port in the operant box. During these sessions, mice received thirty “free” dipper presentations of the reward. We found no significant difference in the percentage of rewards retrieved between WT and *ADGRL3* KO mice (**Fig. S4**).

After successfully learning the task, mice were trained to press a lever using a continuous reinforcement schedule (CRF; **Fig. 5**). We used a fixed-ratio 1 schedule of reinforcement, in which an evaporated milk reward was delivered each time a mouse successfully pressed the lever after its extension into the operant conditioning chamber. The mice were trained over seven sessions to ensure that both WT and *ADGRL3* KO mice successfully completed all trials by pressing the lever and retrieving the reward when it was presented (**Fig. 5B, C**). A significant difference was observed between genotypes in the percentage of lever presses relative to the total number of lever presentations (**Fig. 5B**). *ADGRL3* KO mice acquired lever pressing behavior more quickly than WT mice, achieving approximately 90% completion in the first session compared to ∼22% in controls. To determine whether KO mice associated lever pressing with receiving the evaporated milk reward, we measured the percentage of trials in which mice both pressed the lever and consumed the reward (**Fig. 5C**). In the first session, WT and KO mice had the same percentage of rewarded trials, indicating that despite engaging in more lever pressing, *ADGRL3* KO mice did not exhibit a strong association between lever pressing and reward. In later sessions, *ADGRL3* KO mice learned the lever press-reward association more quickly than WT mice, reaching nearly 100% by the fourth session, whereas WT mice achieved the same level by session seven.

**Figure 5.**
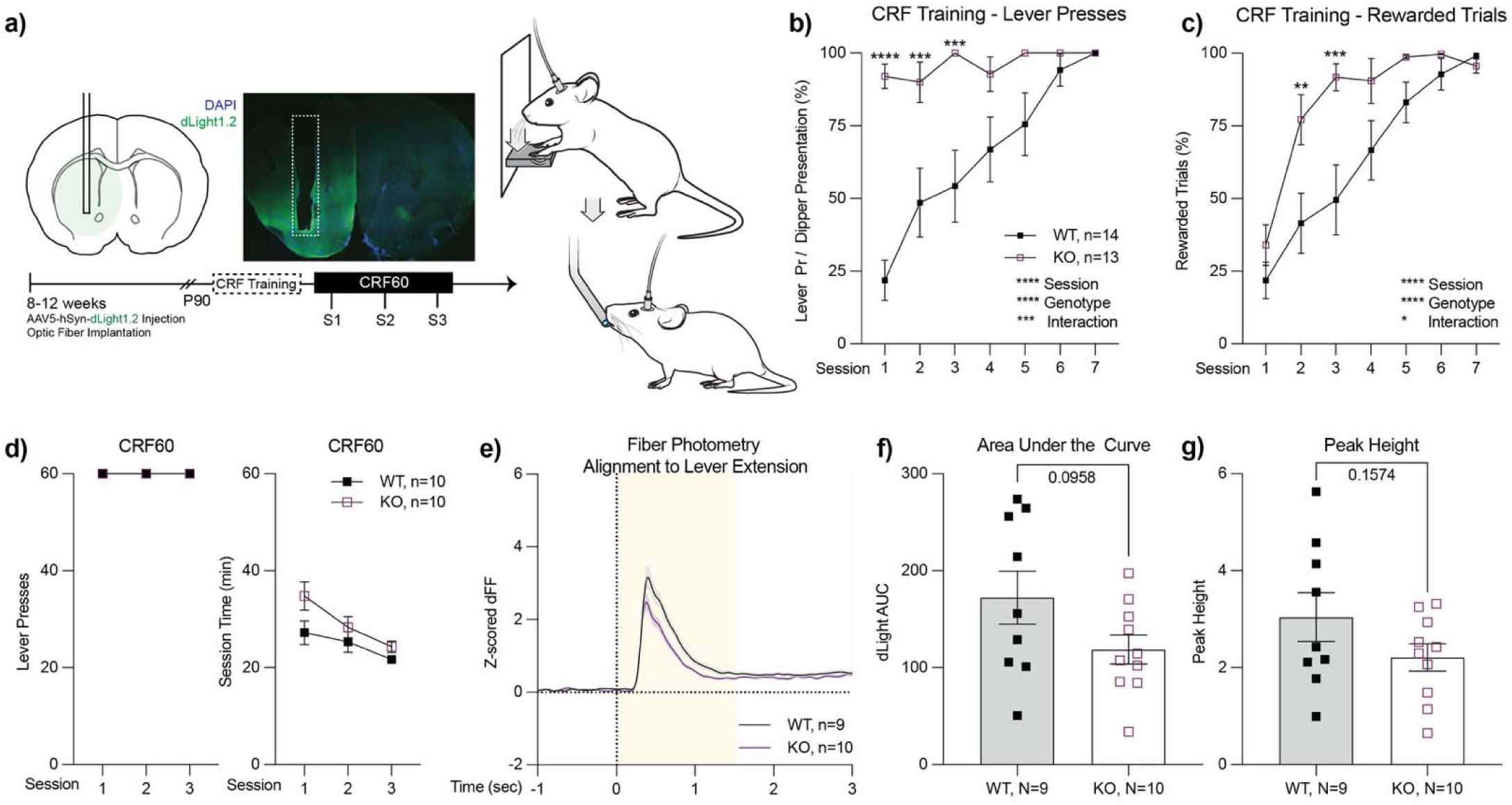
Performance of WT and *ADGRL3* KO mice in a continuous reinforcement task (CRF). **(a)** (Left) Experimental timeline illustrating progression from viral injection and optic fiber implantation to fiber photometry. (Middle) Optic fibers were unilaterally implanted into the nucleus accumbens. A representative coronal section shows dLight1.2 expression in the striatum and the location of the optic fiber, with dashed lines indicating the fiber shaft. (Right) WT and *ADGRL3* knockout (KO) mice were trained on a fixed ratio 1 schedule of reinforcement, where each lever press was rewarded with evaporated milk from a dipper. Training occurred over seven sessions on separate days. **(b)** Percentage of lever presses relative to total possible lever presentations. **(c)** Percentage of trials in which mice lever pressed and consumed the evaporated milk reward. Statistical analysis in **B** and **C** consisted of a two-way ANOVA followed by ídák’s multiple comparison test (*, p<0.05; **, p<0.01; ***, p<0.001; ****, p<0.0001). (N=14 WT, 13 KO). **(d)** Following training, WT and *ADGRL3* KO mice completed a continuous reinforcement task where they could earn up to 60 rewards per session. Dopamine levels were monitored via fiber photometry during this operant task. **(e)** Average dLight1.2 traces aligned to lever extension for WT and *ADGRL3* KO mice. Shaded area (yellow) indicates the timeframe used for AUC calculation. **(f)** Area under the curve (AUC) and **(g)** peak height for the dLight1.2 traces. Statistical analysis consisted of an unpaired t test. (N=9 WT, 10 KO).

To test if *ADGRL3* KO mice exhibited an enhanced learning rate for the CRF task, we analyzed lever press and reward latencies across training sessions (**Fig. S5A, B**). In the first session, *ADGRL3* KO mice pressed the lever significantly faster (42 sec) than WT controls (286 sec) (**Fig. S5A**). However, given the large difference in participation between WT and *ADGRL3* KO mice, we reasoned that comparing numerical sessions might not be appropriate. Thus, we analyzed latency to press during the first session in which each genotype completed at least 50% of trials. Using this approach, we found no significant difference in press latency between genotypes, and both groups showed a decrease in press time across trials within the session. Indeed, by the end of training, both genotypes reached a latency of approximately 5 sec, indicating that, once learned, they could reach a similar performance level.

We applied a similar analysis to reward latency (**Fig. S5B**), where differences between genotypes were less pronounced. Both WT and *ADGRL3* KO mice initially took approximately 3 seconds to retrieve the reward after lever pressing and improved to around 1 second by the end of training. A two-way mixed ANOVA assessing the effects of session and genotype on reward latency revealed that session had the strongest effect. When we analyzed reward latency during the first session in which each genotype completed at least 50% of trials, we found no significant difference between genotypes. Together with the press latency, these findings suggest that once mice achieve similar levels of learned behavior, their response latencies are also comparable.

Following training, WT and *ADGRL3* KO mice were tested in a CRF task where they could earn up to 60 rewards per session. Over the course of three sessions, both genotypes completed all trials and progressively reduced their total session time (**Fig. 5D**). During the task, dopamine release was monitored using *in vivo* fiber photometry with the fluorescent biosensor dLight1.2. The average dLight1.2 signal traces were aligned to lever extension, and the total area under the curve (AUC) was calculated over 0-1.5 seconds to assess potential differences in dopamine release between genotypes (**Fig. 5E, F**). The peak height of the traces was also identified as the maximum change in the signal that exceeded 1 standard deviation above the local baseline (**Fig. 5G**). There were no significant differences between genotypes.

We also investigated the relationship between dopamine release and lever press latency during the task (**Fig. S6**). We found that greater dopamine release was associated with reduced lever press latency. Both WT and *ADGRL3* KO mice displayed this negative correlation, and there was no significant difference in the Pearson correlation coefficients between genotypes.

### The fixed interval task reveals longer latencies and lower levels of dopamine release in *ADGRL3* knockout mice

To further investigate the trends in dopamine release observed during the CRF task, we next challenged the mice with a fixed interval task, where they had to wait for a set period before a lever press would trigger a reward delivery (**Fig. 6A**). Mice were tested across five fixed intervals (2, 4, 8, 12, and 24 seconds), with three sessions per interval. During each trial, the lever was extended, allowing the mouse to interact with it; however, until the designated time interval had elapsed, the lever presses did not activate the dipper response. A successful lever press after the interval resulted in delivery of an evaporated milk reward and lever retraction. The number of lever presses during each interval was recorded.

**Figure 6.**
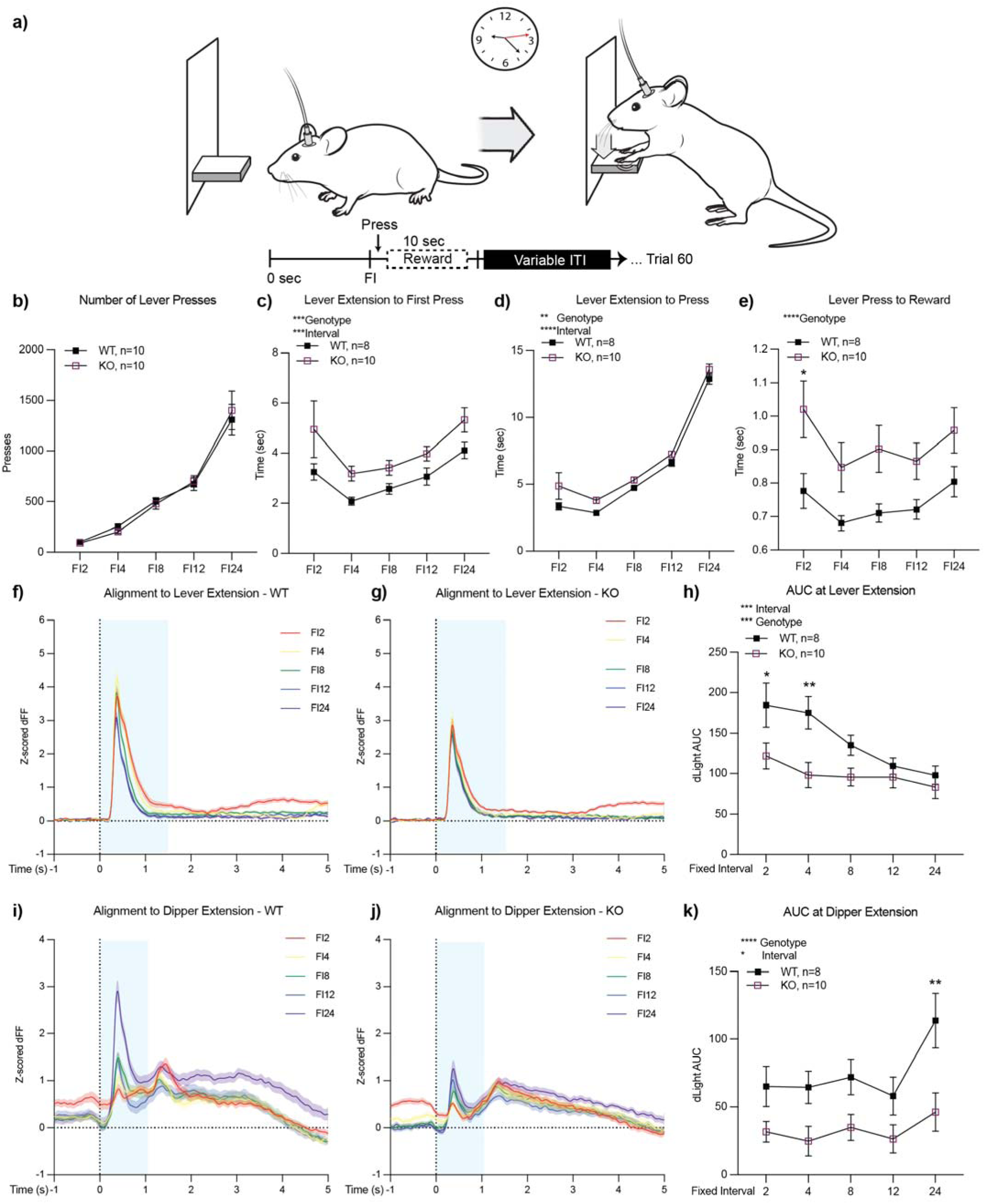
Performance of WT and *ADGRL3* KO mice in a fixed interval task. **(a)** Mice were tested across five fixed intervals (2-24 seconds), with three sessions per interval. During each trial, the lever was extended for the designated interval, after which it remained extended until pressed. A successful press delivered a 10-second evaporated milk reward. Each session included up to 60 rewarded trials, with a variable intertrial interval. **(b)** Total lever presses per interval. No significant genotype differences. (N=10 WT, 10 KO). **(c)** Latency to first lever press, **(d)** average latency to lever press, and **(e)** latency to reward across sessions. All statistical analyses were performed using a two-way ANOVA with ídák’s multiple comparison test (**, p<0.01; ***, p<0.001; ****, p<0.0001). (N=8 WT, 10 KO). **(f, g)** Average dLight1.2 traces aligned to lever extension for WT and *ADGRL3* KO mice. **(h)** Area under the curve (AUC) from 0-1.5 secs. Statistical analysis was performed using a two-way ANOVA with ídák’s multiple comparison test (*, p<0.05; **, p<0.01). (N=8 WT, 10 KO). **(i, j)** Average dLight1.2 traces aligned to dipper extension for WT and *ADGRL3* KO mice. **(k)** AUC from 0-1.0 secs. Statistical analysis was performed using a two-way ANOVA followed by ídák’s multiple comparison test (**, p<0.01). (N=8 WT, 10 KO).

At the behavioral level, there was no difference between genotypes in the total number of lever presses across intervals (**Fig. 6B**). Both WT and *ADGRL3* KO mice increased their number of presses as the fixed interval lengthened. The mice also exhibited a typical “scalloping” pattern within sessions, where their response rates gradually increased as the time for the next reward approached (**Fig. S7**). However, WT mice ramped up lever pressing more rapidly during shorter intervals than *ADGRL3* KO mice, consistent with higher dopamine levels at the time of lever extension. In addition, differences between genotypes were observed when we analyzed latencies to lever press and reward retrieval (**Fig. 6C-E**). A two-way mixed ANOVA revealed significant effects of both interval and genotype on latency to the first lever press. Latency increased with longer intervals, and *ADGRL3* KO mice took significantly longer to initiate a lever press compared to WT controls (**Fig. 6C**). This trend was also observed for average latency to lever press, with KO mice taking longer to press the lever across the session (**Fig. S6D**). The latencies for both genotypes also increased significantly as the interval lengthened. Lastly, latency to reward retrieval was significantly longer for *ADGRL3* KO mice across intervals (**Fig. 6E**).

As with the CRF task, dopamine levels were monitored during the fixed interval task using *in vivo* fiber photometry. We first aligned the dLight1.2 traces to lever extension (**Fig. 6F, G**). We expected dopamine levels to peak during the first fixed interval (2 seconds) and decrease as the interval lengthened, since this decreases the certainty and thereby the expectation of reward. Consistent with this, dLight1.2 AUC decreased with increasing interval lengths (**Fig. 6H**). However, this decrease was less pronounced in *ADGRL3* KO mice, with a delta of 68 units for WT mice and 39 units for KO mice between FI2 and FI24. A two-way repeated measures ANOVA revealed that both interval and genotype significantly affected the AUC at lever extension, with WT dopamine levels being significantly higher than KO mice at shorter intervals.

We next aligned the signal traces to dipper extension (**Fig. 6I, J**), which we defined as the moment the lever retracted and the dipper rose with evaporated milk following a press after the designated interval. With the decay of predictive value by the lever extension, we expected reward prediction to shift to dipper extension, with dopamine levels increasing as the fixed interval lengthened, opposite to the trend observed when aligned to lever extension. In WT controls, this dopamine increase was evident, with a delta of 82 units between FI2 and FI24 (**Fig. 6I**). In contrast, *ADGRL3* KO mice showed a significantly blunted dopamine response to dipper extension, with a delta of only 14.5 units between FI2 and FI24 (**Fig. 6J**). A two-way repeated measures ANOVA confirmed significant effects of both interval and genotype on the AUC at dipper extension, with WT dopamine levels being significantly higher than those of KO mice, particularly at longer intervals (**Fig. 6K**). Peak heights in WT mice (1.1-2.8 units) were also greater than in KO mice (0.7-1.2 units).

### Amphetamine challenge suggests phasic dopamine release capacity is similar between WT and *ADGRL3* KO mice

To better interpret the blunting of *in vivo* dopamine release measured in the behavioral tasks in the *ADGRL3* KO mice, we sought to evaluate whether the dLight1.2 sensor might become saturated, leading to a smaller signal in the context of high tonic dopamine. This was of particular concern given the increased dopamine release observed *ex vivo* in *ADGRL3* KO mice (**Fig. 3**). Therefore, to assess differences in dopamine release capacity between WT and *ADGRL3* KO mice and whether we could observe saturation of the signal, we administered four doses of amphetamine—a drug that induces dopamine release—and monitored dopamine levels using fiber photometry with the dLight1.2 sensor (**Fig. 7**). Concurrently, we assessed locomotor activity in an open field using video tracking and analysis via AnyMaze to capture behavioral correlates of amphetamine response.

**Figure 7.**
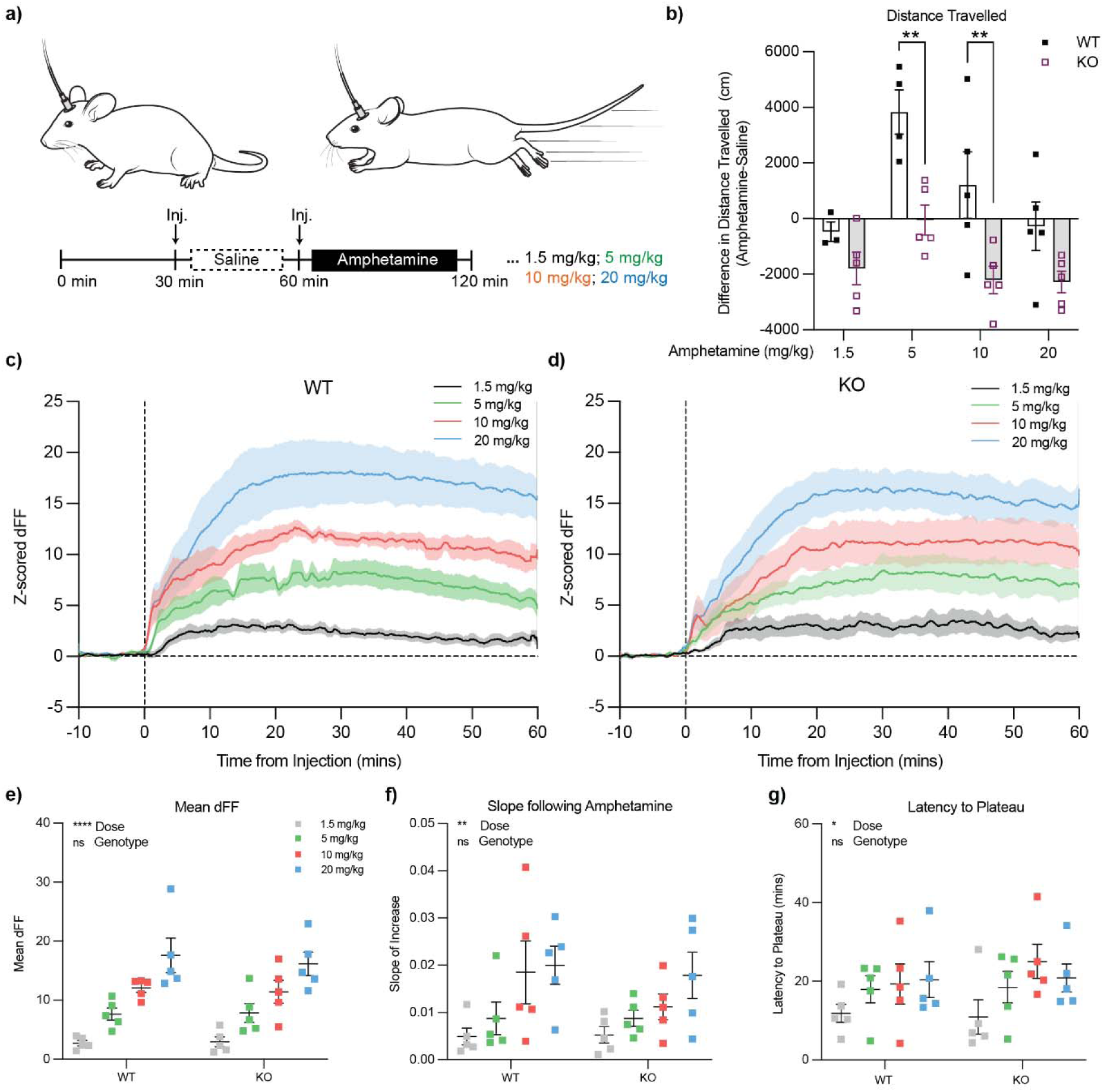
Open field test with amphetamine challenge suggests phasic dopamine release capacity is similar between WT and *ADGRL3* KO mice. (a) Mice injected with AAV5-hSyn-dLight1.2 and implanted with an optic fiber in the nucleus accumbens were used to measure dopamine responses to amphetamine in an open field test. Following a 30-minute habituation period, mice received a saline injection to assess the effects of injection stress on dopamine signaling, followed by amphetamine administration (1.5, 5, 10, or 20 mg/kg subcutaneous) on separate days. Dopamine activity was recorded using fiber photometry, and locomotor behavior was tracked with AnyMaze software (Stoelting Co., Wood Dale, IL). **(b)** Difference in average distance travelled by WT and *ADGRL3* KO mice injected with saline versus varying doses of amphetamine. Statistical analysis was performed using a two-way ANOVA followed by ídák’s multiple comparison test (**, p<0.01). ****, p<0.0001 for genotype and ***, p<0.001 for dose. (N=10 WT, 10 KO). Average dLight1.2 traces aligned to amphetamine injection for **(c)** WT and (d) *ADGRL3* KO mice. **(e)** Mean dF/F, **(f)** slope, and **(g)** latency to reach plateau for WT and *ADGRL3* KO mice following amphetamine injection at the four doses. Statistical analysis was performed using a two-way ANOVA (*, p<0.05; **, p<0.01; ****, p<0.0001). (N=10 WT, 10 KO).

After 30 minutes of habituation in an open field, mice were injected with saline to assess the effects of injection stress on dopamine release, followed by amphetamine administration (1.5, 5, 10, or 20 mg/kg subcutaneous) (**Fig. 7A**). Locomotor activity was continuously monitored throughout the experiment. Amphetamine dose-response tests typically produce a bell-shaped curve of locomotion, with lower doses enhancing locomotor behavior and higher doses inducing stereotypic movements (*26*). In WT controls, both 5 mg/kg and 10 mg/kg amphetamine increased the distance travelled compared to the saline condition, whereas the 20 mg/kg resulted in a reduction of locomotor activity (**Fig. 7B**). In contrast, *ADGRL3* KO mice exhibited reduced locomotor activity across all amphetamine doses.

We then aligned the average dopamine signal traces to amphetamine injection (**Fig. 7C, D)**. Despite the pronounced differences in locomotor behavior between genotypes, there were no statistically significant differences in the mean dFF (**Fig. 7E**), the slope of the curve following amphetamine injection (**Fig. 7F**), or the latency to plateau (**Fig. 7G**) for the dopamine signal traces. We also examined spontaneous dopamine release (**Fig. S8**). The total number of spontaneous events did not differ significantly between WT and *ADGRL3* KO mice (**Fig. S8A, D**). Notably, in both genotypes, spontaneous events decreased with amphetamine treatment, with the greatest reduction observed at the highest dose. Event duration increased in a dose-dependent manner with amphetamine treatment, but there were no genotype differences (**Fig. S8B, E**). The peak height for spontaneous dopamine events remained unchanged by amphetamine treatment and was similar in both WT and *ADGRL3* KO mice (**Fig. S8C, F**). Together, these data suggest that dopamine release capacity is comparable in WT and *ADGRL3* KO mice. The blunted locomotor response in *ADGRL3* KO mice may therefore reflect either increased sensitivity to dopamine leading to rapid onset of stereotypy, or changes in postsynaptic signaling that dampen behavioral output.

## Discussion

In this study, we show that ADGRL3 plays a critical role in modulating dopaminergic function, with potentially distinct effects on dopamine terminals and cell bodies. Rather than uniformly enhancing or suppressing signaling, ADGRL3 influences dopamine release through mechanistically diverse pathways, resulting in both hypo- and hyperdopaminergic phenotypes.

### How is dopamine release altered in *ADGRL3* KO mice?

We first assessed electrically evoked dopamine release *ex vivo* in brain slices using fast-scan cyclic voltammetry (FSCV) (**Fig. 4**). In both the dorsal and ventral striatum of *ADGRL3* KO mice, dopamine release was significantly higher compared to WT controls. In contrast, reuptake in both regions remained unchanged.

We next measured dopamine release in the nucleus accumbens (ventral striatum) of freely behaving mice during a continuous reinforcement and a fixed interval task using *in vivo* fiber photometry with the dLight1.2 biosensor (**Figs. 5, 6**). While there were no significant differences in dopamine release during the continuous reinforcement task, *ADGRL3* KO mice exhibited significantly lower dopamine release during the fixed interval task, where they had to wait for a reward at increasing intervals (2, 4, 8, 12, and 24 seconds). Based on our *ex vivo* slice experiments, we expected that dopamine release in this brain region of *ADGRL3* KO mice would be higher than WT controls (**Fig. 4F-H**). One possible explanation for this discrepancy is that dopamine release in this task is regulated by inputs at the level of the dopamine cell body, while in the voltammetry experiments terminals were stimulated.

We then investigated whether the observed differences in dopamine release during the fixed interval task could be attributed to differences in release capacity, which would, in turn, affect the phasic dopamine response. To assess this, we administered varying doses of amphetamine to the WT and *ADGRL3* KO mice and monitored dopamine release using fiber photometry (**Fig. 7**). If the *ADGRL3* KO mice had an elevated release capacity, we reasoned that their dopamine signal might plateau at lower doses, reaching a ceiling effect sooner than WT mice. That said, the dLight1.2 sensor is reported to produce large responses at low dopamine concentrations (e.g., 100 nM) without approaching saturation (*25*, *27*).

In our data, amphetamine evoked substantially larger fluorescence changes than those observed during behavior, indicating that the sensor operates well below saturation under behavioral conditions (**Fig. 7**). Moreover, we observed comparable, dose-dependent increases in dopamine release across genotypes, and no evidence of a ceiling effect. These findings suggest that the two groups have similar dopamine release capacity and that the differences observed during behavior are accurate and unlikely to reflect sensor limitations.

### Why doesn’t enhanced dopamine release *ex vivo* translate to *in vivo* conditions?

The magnitude, spread, and duration of dopamine release is finely regulated by diverse inputs (*28*). The discrepancies between dopamine release and reuptake in our data suggest that the mechanisms regulating dopamine signaling are context-dependent and differ between *ex vivo* and *in vivo* environments. *Ex vivo* slice preparations lack the physiological tonic and phasic firing of dopamine neurons and are subject to non-specific stimulation, which concurrently activates local cholinergic and other neuromodulatory inputs (*29*, *30*). This can lead to desensitization of nicotinic acetylcholine receptors (*31*, *32*), disrupted D2 autoreceptor feedback (*33–35*), and abnormal calcium dynamics, all of which may emphasize genotype-dependent differences in release and reuptake. In contrast, *in vivo* recordings preserve intact circuit-level regulation, including dynamic modulation by ongoing synaptic input and feedback mechanisms, which may buffer or compensate for differences in dopamine kinetics. Supporting this, a recent study showed that implementing low-frequency tonic stimulation in striatal slices better replicates *in vivo*-like dopamine dynamics, including enhanced sensitivity to D2 autoreceptor regulation (*36*). Their findings underscore how standard slice protocols may fail to capture physiologically relevant presynaptic control, contributing to the divergent outcomes observed between experimental approaches.

### Why is dopamine release reduced in select behavioral contexts?

*ADGRL3* KO mice had reduced dopamine release during the fixed interval task compared to WT controls (**Fig. 6**). In all fiber photometry experiments, we expressed dLight1.2 in the nucleus accumbens, a central component of the ventral striatum that plays a key role in cue-reward association. Based on our *ex vivo* slice experiments (**Fig. 4F-H**), this reduction in dopamine release was unexpected. As noted above, this effect could arise from altered regulation at the level of the dopamine cell bodies rather than at the terminals. This hypothesis is strengthened by our behavioral data for the same task (**Figs. 6B-E, S7**), which show increased latencies to both press and retrieve rewards in *ADGRL3* KO mice. These delays may reflect reduced task engagement, decreased motivation, or a weaker cue-reward association, any of which could lead to reduced dopamine release *in vivo*, even if terminal release capacity remains intact.

Dopaminergic neurons in the midbrain are typically activated by reward-predictive cues (e.g., lever extension), but if upstream neuronal input conveying these cues is diminished, dopamine neuron firing may be blunted. Importantly, our data suggest a behavioral context-dependent reduction in cue-related input, rather than an intrinsic deficit in dopamine neuron function. In contrast, *ex vivo* electrical stimulation bypasses these upstream circuits and directly activates terminals, potentially revealing increased excitability due to local modulation—such as enhanced acetylcholine-evoked dopamine release. This could explain the apparent discrepancy between the *ex vivo* findings and the *in vivo* release patterns observed during behavior.

### How consistent are our findings on dopamine signaling with prior studies?

Our *ex vivo* approach differs from prior studies using an *ADGRL3* KO rat model, where FSCV was used to investigate spontaneous dopamine release in neostriatal slices (*17*). Like our observations with electrically evoked release in the dorsal striatum, *ADGRL3* KO rats exhibited increased spontaneous dopamine release.

However, in contrast to our findings, dopamine reuptake was faster in KO rats than in WT controls, which the authors attributed to increased striatal expression of the dopamine transporter (*16*). We suggest that this discrepancy in reuptake kinetics between our work likely arises from differences in the nature of the dopamine release studied – evoked versus spontaneous. Spontaneous release tends to be slower (*37*, *38*), lasting up to 10 seconds in the referenced study, and more variable, whereas electrically evoked release is rapid and leads to greater dopamine release.

Interestingly, while this manuscript was in preparation, a study using anesthetized *ADGRL3* KO rats assessed stimulation-evoked dopamine release *in vivo* via amperometry (*39*). This technique is similar to our *ex vivo* analyses in that both are electrochemical and provide direct measures of dopamine release. However, unlike slice preparations, *in vivo* amperometry preserves circuit-level connectivity and intact physiological context. The authors assessed dopamine release in two brain regions: the nucleus accumbens core and the medial prefrontal cortex. They reported that *ADGRL3* KO rats had a hypodopaminergic mesocorticolimbic profile, which includes all projections from the ventral tegmental area to regions like the prefrontal cortex and nucleus accumbens. However, their region-specific measurements revealed no significant genotype differences in dopamine release in either brain area. The broader interpretation is nevertheless reminiscent of the reduced cue-induced dopamine release we observed in the nucleus accumbens during the fixed interval task (**Fig. 6**).

The authors of the above study also reported reduced dopamine transporter activity in the prefrontal cortex, but no difference in reuptake kinetics in the nucleus accumbens (*39*). These findings align with our *ex vivo* FSCV and *in vivo* amphetamine challenge data, where dopamine reuptake dynamics in the nucleus accumbens were similar between genotypes (**Fig. 4 Fig. S8**).

### How does knockout of *ADGRL3* influence dopamine-related behaviors?

*ADGRL3* KO mice have exhibited disruptions across multiple dopamine-related behaviors, including motor control, learning, and motivation (*13–15*). Consistent with these reports, we observed similar behavioral phenotypes in our study, most notably hyperactivity – a hallmark feature of the *ADGRL3* KO model (**Figs. 1, 2**). In addition, we found that *ADGRL3* KO mice weighed significantly less than WT controls (**Fig. 1A**). Given that disruptions in dopamine signaling, particularly through D2 dopamine receptors (*40*), have been linked to alterations in energy expenditure and activity, it is possible that the dopaminergic dysregulation resulting from *ADGRL3* knockout contributes to the observed differences in body weight.

Consistent with previous findings, our initial experiments confirmed hyperactivity in a novel environment (**Fig. 1**) and further demonstrated that *ADGRL3* KO mice remain hyperactive in the absence of light as a regulatory cue (**Fig. 2**). Notably, these mice showed altered activity patterns in response to light during the inactive phase, possibly reflecting increased light sensitivity compared to WT controls. However, it is also possible that this responsiveness reflects compensatory rest following elevated activity during the dark phase. The combination of hyperactivity and light-modulated rest may contribute to the observed trend toward higher interdaily stability. Given dopamine’s established role in regulating sleep-wake cycles, future studies could test how constant darkness influences rest-activity rhythms in *ADGRL3* KO mice.

Beyond general hyperactivity, we also explored how altered dopamine dynamics impact goal-directed behavior. *ADGRL3* KO mice had higher response rates in both the gustometer task (**Fig. 3**) and training for the continuous reinforcement task (**Fig. 5B, C**). While this increased engagement may reflect disrupted dopamine signaling affecting goal-directed behavior, it could also be a secondary consequence of the hyperactivity observed in *ADGRL3* KO mice. Additionally, behavioral differences were observed in lever-press and reward-retrieval latencies (**Fig. 6C-E**). Across all measures, *ADGRL3* KO mice exhibited significantly longer latencies, with the most pronounced effect seen in the latency between lever pressing and reward retrieval (**Fig. 6E**). This delay suggests a difficulty in disengaging from lever pressing to consume the reward or decreased cue-induced motivation to respond, both of which could stem from lower dopamine release during the task (**Fig. 6F-K)**. This could also be attributed to behavioral inflexibility secondary to hyperactivity.

It is also important to note that, in addition to the behavioral findings reported above, we encountered significant challenges in maintaining a colony of *ADGRL3* KO mice. Although previous studies have not reported breeding difficulties, we were unable to obtain viable litters from homozygous KO breeding pairs. Female *ADGRL3* KO mice consistently showed poor maternal behavior, often displaying agitation when housed with males and, in many cases, cannibalizing their pups shortly after birth. Due to these issues, we maintained the colony using heterozygous breeding pairs or by pairing a homozygous KO male with a heterozygous female. While maternal behavior is not strictly a result of intact dopamine signaling, the mesolimbic dopamine system is known to play a causal role in maternal motivation (*41*).

## Conclusions

In summary, our findings highlight the complex role of ADGRL3 in modulating dopamine dynamics and behavior. *ADGRL3* KO mice exhibit significant differences in dopamine release, with increased release in *ex vivo* slice preparations but reduced release during specific behavioral tasks *in vivo.* These neurochemical changes are accompanied by an increased instrumental responding and hyperactivity, behaviors consistent with enhanced dopamine release. At the same time, we report increased response latencies during a fixed interval task, suggesting impaired cue-induced motivation consistent with lower dopamine release. These results support a model in which loss of ADGRL3 produces a complex behavioral phenotype by simultaneously enhancing and diminishing dopaminergic function through distinct pre- and postsynaptic mechanisms. This dual effect likely operates across different neural circuits, with behavioral outcomes varying depending on the specific behavioral context.

Importantly, these results point toward broader implications for therapeutic strategies. Mechanisms that control and shape dopamine release at the terminal, especially those involving non-canonical regulators like ADGRL3, may offer more targeted and nuanced intervention points than traditional approaches that directly manipulated dopamine receptors or transporters. Such conventional strategies often act broadly across neural circuits, are prone to receptor desensitization and compensatory adaptations (*42–45*), and are limited by a narrow therapeutic window (*46*, *47*). In contrast, targeting upstream modulators like ADGRL3 could allow for circuit-specific modulation with potentially fewer side effects. While further research will be needed to unravel the specific signaling mechanisms by which ADGRL3 modulates dopamine in the striatum, our findings support that ADGRL3 may emerge as a novel therapeutic target for disorders involving dopamine dysfunction.

## Materials and Methods

### Mice

Adult male and female mice (129S4/SvJae and C57 mix)(*15*) null for *ADGRL3* (MGI:4973916) were housed in groups of five per cage in a 12-hour light/dark cycle (6:00 – 18:00) at 22°C. Food and water were provided *ad libitum*. This mouse line contains a genomic insertion that disrupts the mucin stalk domain and is predicted to eliminate any intracellular signaling from ADGRL3, if not all functions. Genetic monitoring (Transnetyx miniMUGA) confirmed that experimental mice were inbred and of excellent genotyping quality, with mixed background primarily comprising C57BL/6J, C57BL/6NCrl, and some 129S4/SvJaeJ ancestry. All experimental procedures were conducted in accordance with NIH guidelines and were approved by the Institutional Animal Care and Use Committees of Columbia University and the New York State Psychiatric Institute.

### Open Field

Locomotor activity in wild-type and *ADGRL3* knockout mice was assessed from 6 to 12 weeks of age. Mice were placed in a plexiglass enclosure equipped with infrared photobeams measuring 42×42×38 cm for 60 mins undisturbed (Med Associates, St. Albans, VT). Activity was recorded using Motor Monitor software (Kinder Scientific, Poway, CA), which detects infrared beam breaks within the enclosure. The software tracked horizontal movements, rearing instances, and time spent in the center and periphery of the enclosure, defined as a 21 × 21 cm grid at the center. Basic movements were measured in counts and defined as discrete episodes of whole-body movement reflecting overall activity levels. Rearing behavior, also measured in counts, was defined as vertical exploratory behavior (i.e., standing on hind limbs). Data for each variable were analyzed by summing measurements over the entire 60-min period.

### Piezo Sleep Study

The Piezo-Sleep mouse behavioral tracking system (Signal Solutions LLC, Lexington, KY) was used to monitor activity levels and measure sleep/wake cycles in mice using established protocols(*48*, *49*). Over a period of five consecutive days, mice were individually housed in chambers equipped with sawdust bedding, with *ad libitum* access to food and water, under a 12-hour light/dark cycle (6:00-18:00). The system’s floor sensors tracked sleep bouts, respiration, temperature, and movement. Activity counts (scaled between 0.0-0.3) represent normalized movement intensity per ten minutes, while the activity amplitude represent the intensity of detected movements during active periods. Interdaily stability and intradaily variability were calculated as outlined previously (*50*).

### Gustometer

The day before gustometer training, mice were switched to a 2% w/v citric acid solution in their home cage water bottles. This approach allows mice to regulate their own water intake but imposes a mild restriction due to the aversive taste of citric acid. Mice were weighed daily and observed for normal behavior. On the first day of training, 2–3 mice were placed in the “Davis Rig” gustometer (Med-Associates, St. Albans, VT) for 30 minutes to learn licking behavior at the lick-port. During this session, the sipper was made available for 5–15 seconds, followed by a fixed intertrial interval. On the second day, each mouse underwent individual training with two sipper tubes for 30 minutes, following a similar procedure. On the testing day, mice were introduced to a 6-bottle choice paradigm, offering evaporated milk concentrations ranging from 0% to 100%. The task lasted 30 minutes, with a 20-second intertrial interval. Sampling from the sipper tubes followed a pseudorandom order.

### Fast-scan cyclic voltammetry in brain slice

Male and female mice (3-6 months old) were euthanized by cervical dislocation. The brains were then quickly removed and immersed in an ice-cold sucrose cutting solution (10 mM NaCl, 2.5 mM KCl, 25 mM NaHCO_3_, 0.5 mM CaCl_2_, 7 mM MgCl_2_·6H_2_O, 1.25 mM NaH_2_PO_4_·6H_2_O, 180 mM sucrose, and 10 mM glucose), which was oxygenated with a 95% O_2_/5% CO_2_ mixture and adjusted to a pH of 7.4. Coronal sections that included the striatum were cut at a thickness of 250 µm, using a slicing speed of 0.24 mm/s and an amplitude of 1.7 mm. Each brain section was allowed to recover for 30 minutes at 34°C in artificial cerebrospinal fluid (ACSF; 125 mM NaCl, 2.5 mM KCl, 25 mM NaHCO_3_, 1.5 mM CaCl_2_, 1 mM MgCl_2_·6H_2_O, 1.25 mM NaH_2_PO_4_·6H_2_O, and 10 mM glucose). Slices were then bubbled in oxygenated ACSF at room temperature for at least 30 min prior to transfer to a recording chamber.

Fast-scan cyclic voltammetry (FSCV) recordings of evoked dopamine release were conducted following protocols outlined in previous studies (*33*, *36*). Striatal slices were placed in a recording chamber, secured with a slice anchor, and perfused with ACSF at a constant rate of 2 mL/min at 34°C. Recording time did not exceed 3 hours. Dorsal and ventral striatum recordings were made from each slice, with the order of testing between brain regions randomized.

To prepare for recording, a sharpened bipolar concentric electrode (400 µm maximum outer diameter; Pt/Ir; WPI) was positioned in either the dorsal or ventral striatum of each brain slice. A carbon fiber electrode (5 µm diameter, cut to a length of 100-200 µm) was then placed approximately 150 µm from the stimulation electrode at a depth of about 50 µm. A triangular voltage waveform, moving from −450 mV to +800 mV over 8.5 ms (a ramp rate of 294 V/s), was applied to the carbon fiber electrode every 100 ms. The resulting current was recorded using an Axopatch 200B amplifier (Molecular Devices, Foster City, CA), filtered with a 10 kHz low-pass Bessel filter, and digitized at 25 kHz (ITC-18 board, InstruTECH, Great Neck, NY). FSCV data acquisition and analysis were performed using custom in-house procedures developed in Igor Pro (WaveMetrics Inc., Lake Oswego, OR).

During recording, slices were stimulated with the bipolar concentric electrode using an Iso-Flex stimulus isolator (AMPI), which was triggered by TTL pulses generated using a custom-built Igor Pro routine. Single pulses (0.1-1.0 ms, 200 µA) were delivered every 2 minutes for a total of 10 minutes, and the three middle peaks were analyzed. Carbon fiber electrodes were calibrated by quantifying background-subtracted voltammograms in standard solutions of 1 µM dopamine prepared in 0.1% HCl.

### Stereotaxic Surgery and Optic Fiber Implantation

Mice (≥8 weeks old) were initially anesthetized with 3% isoflurane at a flow rate of 500 mL/min, then maintained at 1-2% isoflurane at 100 mL/min throughout the procedure. Using a Hamilton syringe (Model 701) and a programmable nanoliter injector, 375 nL of AAV5-hSyn-dLight1.2 (Addgene Cat. #111068-AAV5) was unilaterally injected into the nucleus accumbens.

Stereotaxic injections were guided by bregma-based coordinates: AP, +1.7 mm; ML ±1.2 mm; DV, −4.2, −4.1, and −4.0 mm (125 nL per DV site). Injections were spaced 1 minute apart, with a 5-minute pause following the final injection to allow the virus to diffuse into the tissue.

Following the virus injection, 400 µm fiber optic cannulas (Doric Lenses Inc., 5.0 mm fiber length, FLT tip) were carefully lowered to a depth of −4.0 mm and secured to the skull using dental cement (C&B Metabond Quick Adhesive Cement System) and machine mini screws. A subcutaneous injection of 500-1,000 µL saline was then administered, and the mouse was allowed to recover on a heating pad for at least 15 minutes. Mice were monitored for a minimum of 3 days post-surgery and received subcutaneous carprofen (5 mg/mL) as needed for pain management. Mice with implanted cannulas began behavioral training approximately 4 weeks after surgery. At the end of the experimental period, the mice were perfused with 4% paraformaldehyde. Following perfusion, their brains were processed to verify the expression of the virus and the location of the optic fibers.

### Operant Apparatus

All operant behavioral tasks (CRF and FI) were conducted in operant chambers (Med-Associates, St. Albans, VT) housed within sound-attenuating cubicles. Each chamber was equipped with retractable levers, a liquid dipper, and an LED house light that illuminated the chamber during tasks. The retractable levers were positioned 5 cm away from the feeder trough on the same wall. Head entries into the reward magazine were detected using an infrared photocell detector, and each dipper delivered a drop of evaporated milk, approximately 15 µL in volume. The Med-PC Behavioral Control Software Suite was used to control each operant chamber and record the acquired data.

Adult mice (>90 days) were first trained to access the dipper, followed by lever press using a continuous reinforcement schedule. After completing operant training, the mice underwent fixed interval training. Mice were weighed daily, and food restricted to 80-90% of their baseline weight, while access to water remained unrestricted.

### In vivo fiber photometry

The setup and operation of the fiber photometry equipment were described previously (*51*). A 4-channel programmable LED Driver (Doric Lenses, Quebec, Canada) was connected to two sets of connectorized LEDs, one at the isosbestic signal 405 nm and the other at 465 nm. The 405 nm LEDs were routed through 405-410 bandpass filters, while the 465 nm LEDs passed through a 460-490 nm GFP excitation filter using two 6-port Doric fluorescent minicubes. Excitation LEDs were sinusoidally modulated using Synapse software and RZ5P Multi I/O Fiber Photometry Processors (Tucker-Davis Technologies, Alachua, FL) at 210 Hz (405 nm) and 330 Hz (465 nm). The powers at the patch cord for 405 nm and 465 nm LEDs were set to 6-12 µW or 30 µW, respectively. Fluorescence emissions from the implanted optic fiber cannula were transmitted via a fiber-optic patch cord (400 µm/0.48 NA) connected to the minicubes, then filtered through 500-540 nm GFP emission filters, and directed to photodetectors with the gain set to “DC low.” Fluorescent signals were digitized at 1017 Hz and demodulated into isosbestic and GFP components with a 6 Hz low-pass filter.

Fiber photometry data were analyzed using custom MATLAB scripts. Signals were downsampled by a factor of 10 before further analysis. The 405 nm channel served as an isosbestic control to correct for movement artifacts and photobleaching, while the 465 nm channel captured fluorescence changes from dLight1.2, reflecting dopamine binding. To calculate the change in fluorescence (dFF), a fitted 405 nm signal was generated using a least-squared linear regression of the 405 nm signal against the 465 nm signal.This fitted control was then used to normalize the 465 nm signal using the follow formula:

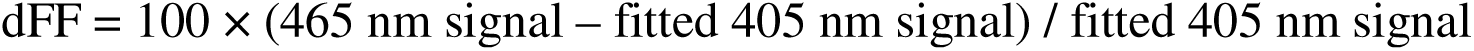

To account for variability in dLight1.2 expression levels or fiber placement relative to the source of fluorescence, the resulting dFF signal was z-scored.

For each session, z-scored dFF data were aligned to key behavioral events such as lever extension, dipper presentation, or reward delivery, on a trial-by-trial basis. A baseline was defined as the average signal during the inter-trial interval (ITI, −5 to −0.5 seconds before the first behaviorally relevant event of the trial) and was subtracted from the aligned data. Baseline-corrected dFF traces were then averaged across trials for each session and subsequently across session for each mouse. Peak amplitude and area under the curve (AUC) were calculated from the behaviorally aligned, z-scored dFF signal within specific time windows following each event: 0-1.5 sec after lever extension, 0-1 sec after dipper presentation, and 0-3 sec after reward delivery.

Spontaneous events were identified from the z-scored dFF during ITIs. The MATLAB function “findpeaks” was used to detect within ITIs for each session. To ensure relevance, only peaks with a prominence exceeding two standard deviations above the session mean were included. For each detected peak, a 4-second window (−2 to +2 seconds relative to the peak) centered on the detected peak was extracted, and the prominence, peak amplitude, and width were calculated.

### Dipper Training

Mice completed four sessions of dipper training, during which they received “free” dipper presentations without needing to press a lever. Each time the dipper was activated, it stayed in the port for 10 seconds before retracting. Head entries and poke latencies were recorded. For the first session, mice received 30 dipper presentations, each separated by a variable inter-trial interval (ITI) of 30 seconds. For the remaining sessions, mice received 60 dipper presentations.

### Continuous Reinforcement Schedule

During lever press training, each lever press was reinforced using a continuous reinforcement (CRF) schedule. After each reinforcement, the levers were retracted and reintroduced following a variable ITI, with ITIs randomly generated to average 20 seconds. The reinforcement involved raising the dipper containing an evaporated milk reward for 5 seconds. Initially, the mice were trained in sessions that ended after the mouse received 30 reinforcements or after 30 minutes, whichever occurred first.

Subsequently, they were trained in sessions that ended after 60 reinforcements or after one hour, whichever occurred first. Training sessions were repeated daily until the mice consistently earned 60 reinforcements.

### Fixed Interval Task

After completing CRF training, mice were tested on a fixed interval (FI) schedule. In these sessions, mice could receive up to 60 reinforcements from pressing the right lever, or the session would end after one hour, whichever came first. After each reinforcement, the lever was retracted and reintroduced following a variable ITI, with the ITIs randomly generated to average 12 seconds. Reinforcement consisted of raising the dipper containing evaporated milk for 10 seconds.

Mice were tested across five fixed intervals: 2, 4, 8, 12, and 24 seconds. Each interval was repeated for three sessions. During these intervals, the lever was extended for the designated fixed time, allowing the mouse to interact with it. The number of lever presses during each interval was recorded, though the lever would only retract once the fixed time had elapsed.

### Open field test with amphetamine challenge and fiber photometry

Select mice that had undergone AAV5-hSyn-dLight1.2 virus injection and fiber optic implantation in the nucleus accumbens were used to detect dopamine changes in response to amphetamine challenge in an open field test. The purpose of this experiment was to determine whether baseline dopamine levels differed between WT and *ADGRL3* KO mice. Four doses of amphetamine (1.5, 5, 10, and 20 mg/kg in 0.9% saline) were administered subcutaneously (s.c.) on separate days, following the behavioral protocol outlined below. This analysis was based off a recent study that compared microdialysis and fiber photometry for measurements of extracellular dopamine (*52*).

On the day of the experiment, mice were weighed, tethered to an optical cable for fiber photometry measurements, and placed in a plastic enclosure measuring 24×45×20 cm with approximately 0.5 cm of bedding. A 30 min habituation period in the open arena was conducted, during which video recording was initiated using AnyMaze software (Stoelting Co., Wood Dale, IL), and fiber photometry signals were sampled as described above (see ***In vivo fiber photometry***). After the habituation period, mice were s.c. injected with 0.9% saline and monitored for 30 minutes to assess whether the injection stress affected dopamine recordings. Finally, the mice were s.c. injected with amphetamine and monitored for an hour.

### Histology

Adult mice were anesthetized with an intraperitoneal (i.p.) injection of 100 mg/kg ketamine and 10 mg/kg xylazine, then perfused with 4% paraformaldehyde (PFA) in phosphate-buffered saline (PBS). After perfusion, the brains were removed and post-fixed overnight in 4% PFA, then transferred to a 0.1% sodium azide solution in PBS. For sectioning, 50 µm slices were cut using a vibratome (Leica, Buffalo Grove, IL, USA) at an amplitude of 1.0 mm and a cutting speed of 1.0 mm/s. The slices were stained overnight at 4°C with a primary antibody against dLight1.2 (chicken anti-GFP; Abcam, Cambridge, UK, ab13970, 1:1,000) prepared in blocking buffer (5% donkey serum, 0.5% bovine serum albumin, 0.1% PBS-Triton). After primary staining, the slices were incubated with an Alexa Fluor-conjugated secondary antibody (goat anti-chicken IgY Alexa Fluor 488; Thermofisher, A11039, 1:500) and DAPI (Thermofisher Scientific, D1306, 1:1,000) for one hour at room temperature in the dark. The stained slices were rinsed in 50 mM Tris (pH 7.5) to remove salts and mounted on glass slides using ProLong Glass Antifade Mountant (ThermoFisher Scientific, Waltham, MA). Viral expression and recording site lesions were confirmed through GFP staining using an SP8 confocal microscope (Leica Microsystems, Wetzlar, Germany).

### Statistical Analysis

Statistical analyses are detailed in the corresponding figure legends. All analyses were performed using GraphPad Prism version 10.4.2 for Mac OS X (GraphPad Software, Boston, MA; www.graphpad.com) or custom scripts written in MATLAB.

## Acknowledgements

The authors thank Eleanor Simpson for valuable discussions regarding data analysis of fiber photometry recordings. We are also grateful to Kirill Martemyanov for providing breeders for the *ADGRL3* knockout mouse line. Illustrations used in this manuscript were sourced from **Sci-Draw**, with thanks to Ethan Tyler and Lex Kravitz (DOIs: 10.5281/zenodo.3925984, 10.5281/zenodo.3925948, 10.5281/zenodo.3925972, and 10.5281/zenodo.3926056), and Roberta Schellino (DOI: 10.5281/zenodo.4966122). We acknowledge Sci-Draw for making this valuable resource openly available.

## Funding

National Institutes of Health grant T32 MH015144 (N.A.P.H.)

Columbia University Institute for Developmental Sciences Award (N.A.P.H.) National Institutes of Health grants MH054137 (J.A.J.) and MH124858 (C.K.) Hope for Depression Research Foundation (J.A.J.)

Miriam’s Magical Memorial Mission (J.A.J.)

## Author contributions (CRediT)

Conceptualization: NPH, CK, JJ Methodology: NPH, ATH, SB, DL, MJ

Formal Analysis: NPH, ATH, SB, EAM, EVM Investigation: NPH, SB, EAM

Data Curation: ATH

Writing – Original Draft: NPH (lead), ATH, CK, JJ

Writing – Review and Editing: NPH (lead), ATH, SB, EAM, DL, MJ, CD, DS, EVM, CK, JJ

Visualization: NPH, ATH

Resources: CD, DS, CK, JJ

Supervision: EVM, CK, JJ Funding Acquisition: CK, JJ

## Data and materials availability

All data needed to evaluate the conclusions in the paper are present in the paper and/or the Supplementary Materials. Data will be shared upon request. Contact the corresponding authors here

