## Supplementary Files for "Altered striatal dopamine regulation in *ADGRL3* knockout mice"

Nicole Perry-Hauser *et al.*

**This PDF file includes:**

Figs. S1 to S8

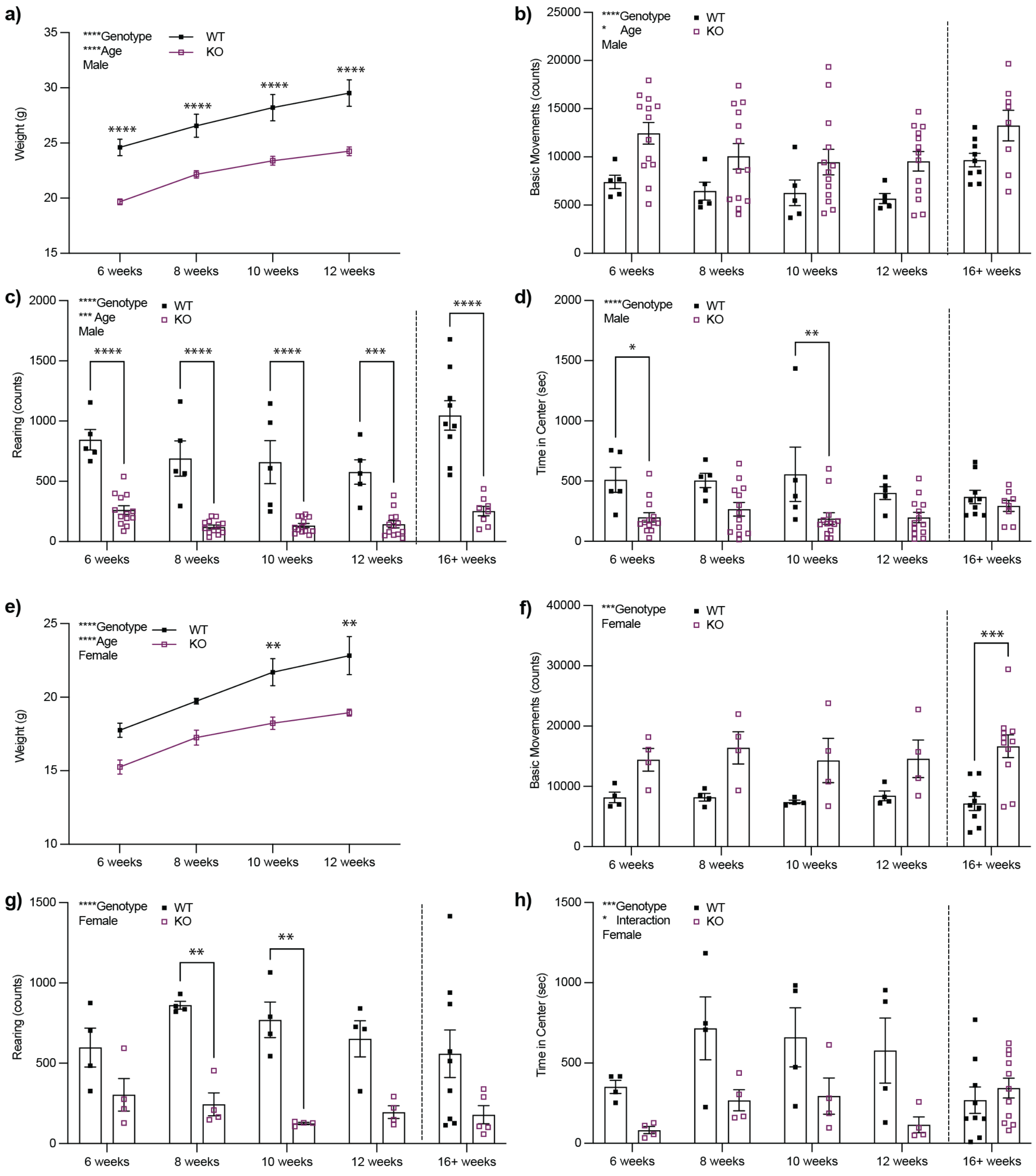


**Fig. S1. *ADGRL3* knockout mice show increased horizontal locomotion and decreased rearing activity across development. (a)** Weight of wild-type and *ADGRL3* knockout male mice (N=5 male WT, 13 male KO at 6-12 weeks of age; N=9 male WT, 8 male KO at 16+ weeks of age). **(b)** Horizontal locomotor activity, **(c)** rearing activity, and **(d)** time in center for WT and *ADGRL3* KO male mice across development. **(e)** Weight of wild-type and *ADGRL3* knockout female mice (N=4 female WT, 4 female KO at 6-12 weeks of age; N=10 female WT, 11 female KO at 16+ weeks of age). **(f)** Horizontal locomotor activity, **(g)** rearing activity, and **(h)** time in center for WT and *ADGRL3* KO female mice across development. A two-way repeated measures ANOVA was performed to analyze the effect of age and genotype on each dependent variable (weight, distance traveled, rearing, or time in center). This was followed by Śídák's multiple comparisons test (*, $p < 0.05$; **, $p<0.01$; ***, $p<0.001$; ****, $p < 0.0001$).

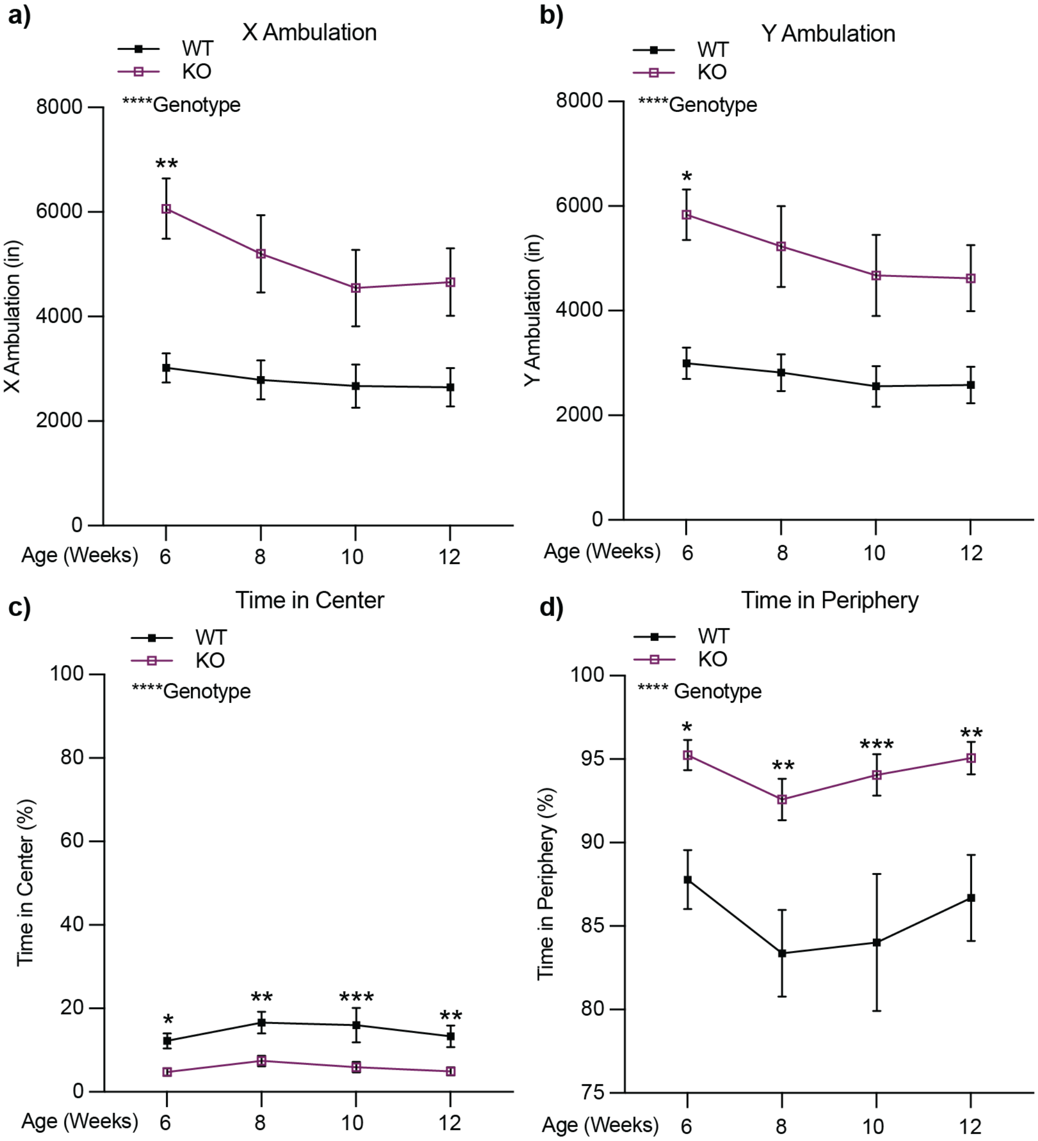


**Fig. S2. *ADGRL3* knockout mice show increased X-Y ambulation and decreased time in center across development.** Wild-type and *ADGRL3* knockout mice (N=9 WT, 17 KO) **(a)** X ambulation (in inches) **(b)** Y ambulation (in inches) **(c)** time in center (%), and **(d)** time in periphery (%) across development. A two-way repeated measures ANOVA was performed to analyze the effect of age and genotype on each dependent variable (distance or percentage of time). This was followed by Śídák's multiple comparisons test (*, $p < 0.05$; **, $p<0.01$; ***, $p<0.001$; ****, $p < 0.0001$).

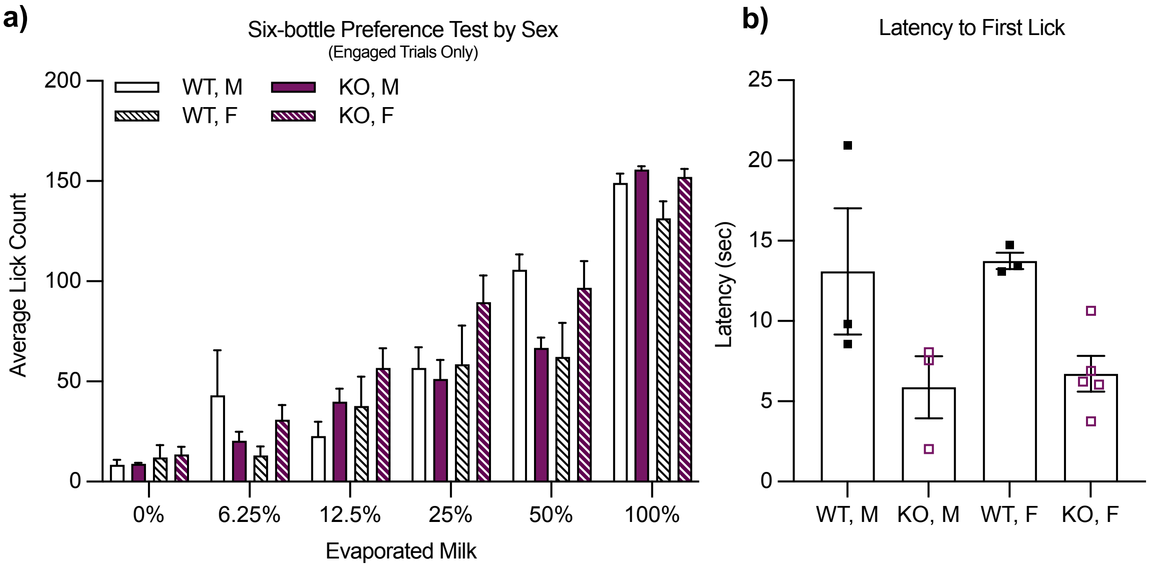


**Fig. S3.** ***ADGRL3*** **knockout mice have similar preference for an evaporated milk reward in a six-bottle test compared to WT mice. (a)** Average licks divided by genotype and sex for 6 different concentrations of evaporated milk (0%, 6.25%, 12.5%, 25%, 50%, and 100%). Statistical analysis consisted of a Three-way ANOVA (****, p<0.0001 for % of evaporated milk). No statistical difference was detected for genotype or sex (N=3 WT female, 3 WT male; 5 KO female, 3 KO male). **(b)** Average latency until first lick by sex. Statistical analysis consisted of an ordinary one-way ANOVA (*, p<0.1).

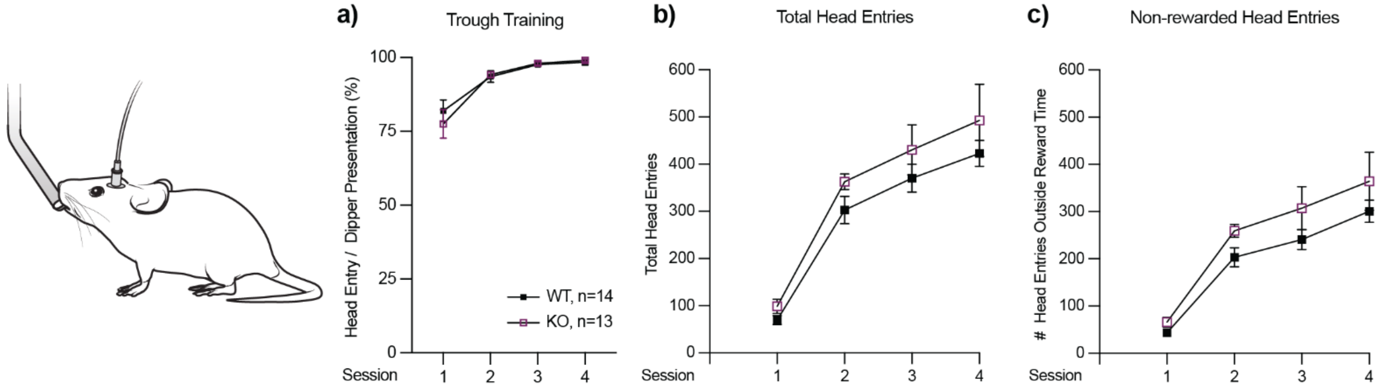


**Fig. S4. *ADGRL3* knockout mice exhibit similar reward retrieval learning as WT mice. (a)** Percentage of head entries relative to total possible dipper presentations. *ADGRL3* KO mice learn to retrieve an evaporated milk reward from an extended dipper at a comparable rate to WT mice. Statistical analysis was performed using a two-way ANOVA followed by Śídák's multiple comparison test (****, $p<0.0001$ for session). (N=14 WT, 13 KO). **(b)** Total number of head entries into the dipper port across sessions. **(c)** Number of head entries recorded outside of reward presentation periods.

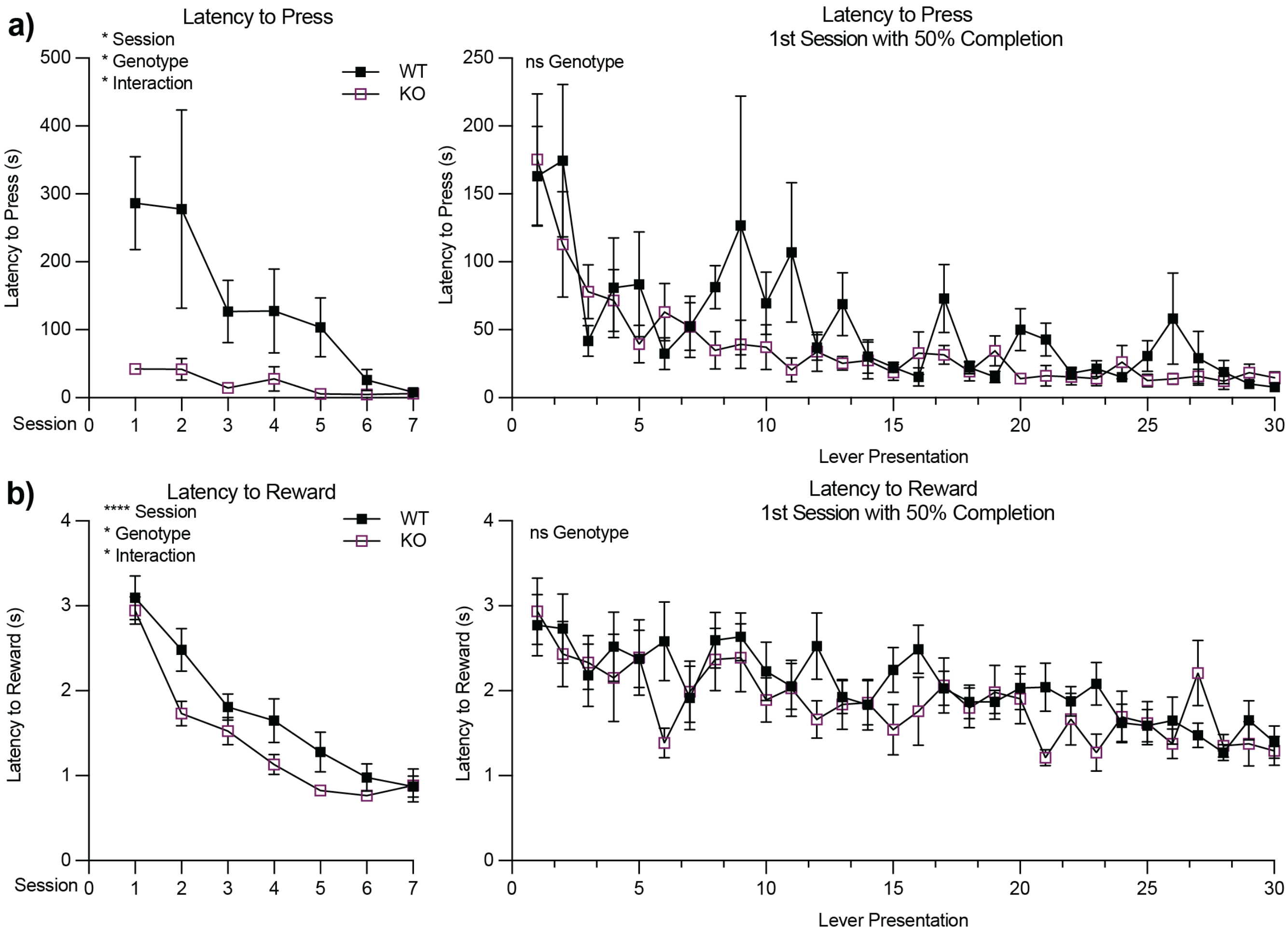


**Fig. S5. Latencies to lever press and reward retrieval during continuous reinforcement training. (a) (Left)** Latency to lever press (sec) across sessions for WT and *ADGRL3* KO mice. Statistical analysis was performed using a mixed effects analysis followed by Dunnett's multiple comparison test (*, p<0.05). (N=14 WT, 13 KO). **(Right)** Latency to lever press across trials during the first session with 50% of the trials completed. No significant difference in latency to lever press was observed under this criterion. **(b) (Left)** Latency to reward (sec) across sessions for WT and *ADGRL3* KO mice. Statistical analysis was performed using a mixed effects analysis followed by Dunnett's multiple comparison test (*, p<0.05; ****, p<0.0001). (N=14 WT, 13 KO). **(Right)** Latency to reward across trials during the first session with 50% of the trials completed. No significant difference in reward retrieval was observed under this criterion.

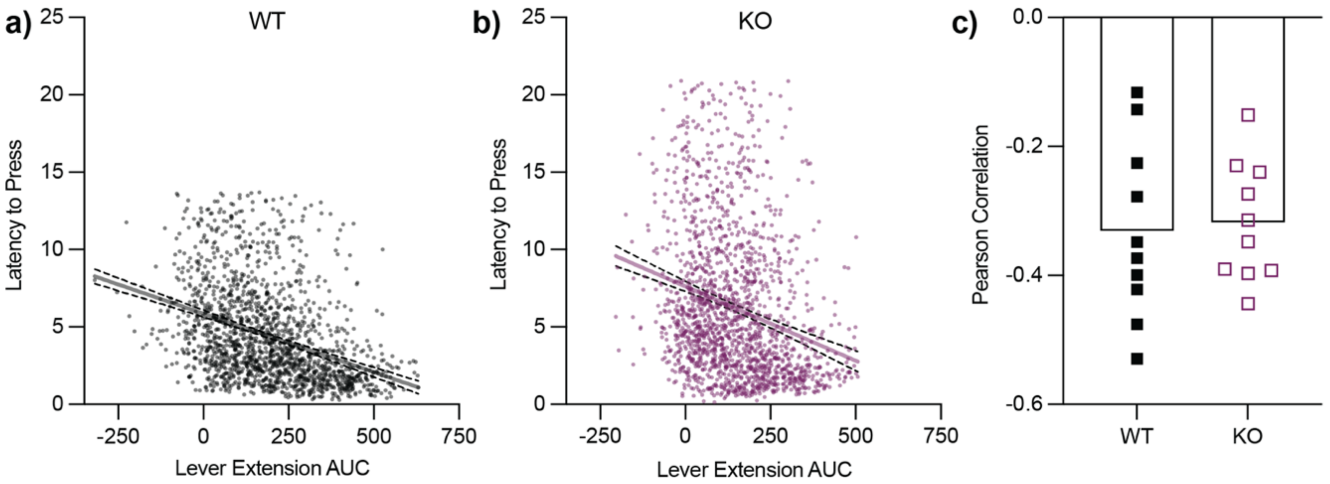


**Fig. S6. Relationship between dopamine release and lever press latency during the continuous reinforcement task.** Correlation between area under the curve and latency to lever press for **(a)** WT and **(b)** *ADGRL3* KO mice during the continuous reinforcement task (CRF60). Both genotypes exhibited a negative correlation, indicating that greater dopamine release was associated with reduced lever press latency. **(c)** No significant difference in Pearson correlation coefficients between WT and *ADGRL3* KO mice.

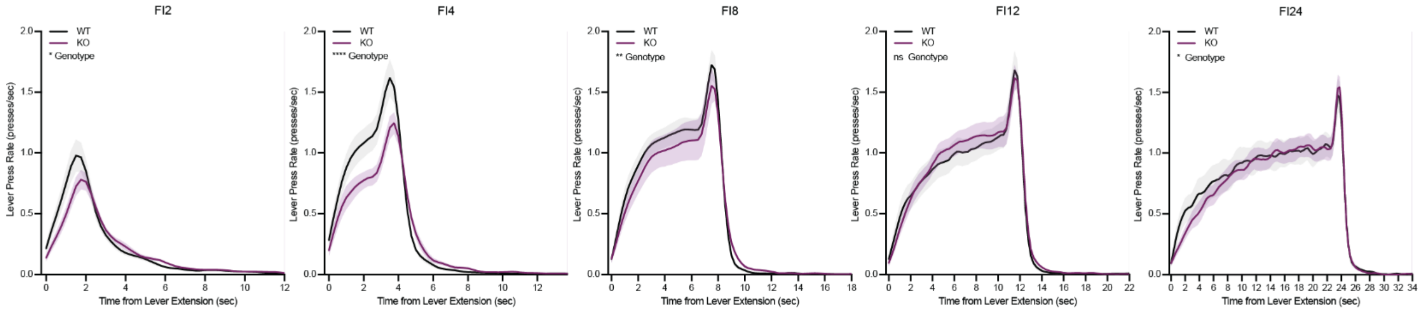


**Fig. S7. Lever press rates increased as the time for the next fixed interval approached.** Lever press rates (presses per second) were plotted across all fixed intervals following the intial lever extension. A characteristic "scalloping" pattern emerged, marked by an initial pause after lever extension, followed by a gradual increase in lever pressing as the fixed interval progressed. Statistical analysis was performed using a two-way ANOVA to analyze the effect of genotype and time on lever press rate (* p<0.05; **, p<0.01; ***, p<0.001; ns, not significant). (N=9 WT, 10 KO).

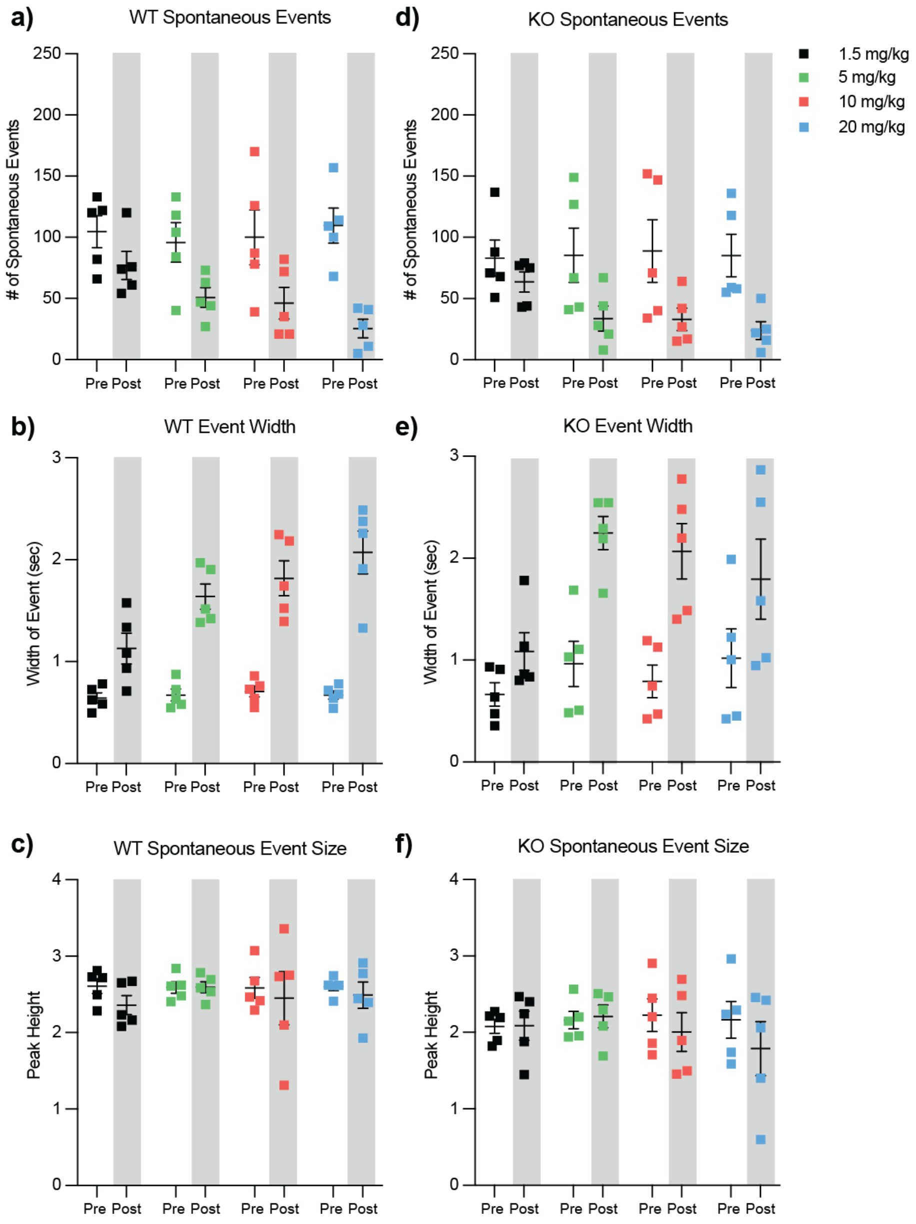


**Fig. S8. Spontaneous photometry events during the open field test with amphetamine challenge showed no significant differences between WT and *ADGRL3* KO mice. (a)** Total number of spontaneous events, **(b)** event duration, and **(c)** peak height for WT mice before and after amphetamine injection. **(d)** Total number of spontaneous events, **(e)** event duration, and **(f)** peak height for KO mice before and after amphetamine injection. Statistical analysis was performed using a two-way ANOVA (*, $p<0.05$; **, $p<0.01$; ****, $p<0.0001$). (N=10 WT, 10 KO).
